# A Distance Method to Reconstruct Species Trees In the Presence of Gene Flow

**DOI:** 10.1101/007955

**Authors:** Lingfei Cui, Laura S. Kubatko

## Abstract

One of the central tasks in evolutionary biology is to reconstruct the evolutionary relationships among species from sequence data, particularly from multilocus data. In the last ten years, many methods have been proposed to use the variance in the gene histories to estimate species trees by explicitly modeling deep coalescence. However, gene flow, another process that may produce gene history variance, has been less studied. In this paper, we propose a simple yet innovative method for species trees estimation in the presence of gene flow. Our method, called STEST (Species Tree Estimation from Speciation Times), constructs species tree estimates from pairwise speciation time or species divergence time estimates. By using methods that estimate speciation times in the presence of gene flow, (for example, M1 (Yang 2010) or SIM3s (Zhu and Yang 2012)), STEST is able to estimate species trees from data subject to gene flow. We develop two methods, called STEST (M1) and STEST (SIM3s), for this purpose. Additionally, we consider the method STEST (M0), which instead uses the M0 method (Yang 2002), a coalescent-based method that does not assume gene flow, to estimate speciation times. It is therefore devised to estimate species trees in the absence of gene flow. Our simulation studies show that STEST (M0) outperforms STEST(M1), STEST (SIM3s) and STEM in terms of estimation accuracy and outperfroms *BEAST in terms of running time when the degree of gene flow is small. STEST (M1) outperforms STEST (M0), STEST (SIM3s), STEM and *BEAST in term of estimation accuracy when the degree of gene flow is large. An empirical data set analyzed by these methods gives species tree estimates that are consistent with the previous results.

## Introduction

Species tree estimation is one of the most fundamental problems in evolutionary biology. Gene trees estimated from sequences sampled from the corresponding species were once treated as the species tree estimate before adavances in sequencing techniques made multilocus data available for routine phylogenetic analysis. Now, it is well-appreciated that an embedded gene tree may not match its underlying species tree, i.e., a gene may have a different evolutionary history from its underlying species (Fitch 1970; Tajima 1983; Pamilo and Nei 1988; Felsenstein 2004). The causes for such incongruence include deep coalescence, horizontal gene transfer (HGT) or lateral gene transfer (LGT), and gene duplication/loss (see Fig. 1). Deep coalescence, also called incomplete lineage sorting (ILS), refers to the case when the coalescent time is deeper into the past than the previous speciation time (Fig. 1a), which might result from large population sizes or short speciation times (Maddison 1997). HGT or LGT refers to gene flow between organisms that is not through reproduction (Fig. 1b). The probability of HGT between distinct species is different. It is widely accepted that in prokaryotes, HGT happens frequently, thus playing an important role in evolution (Boto 2010). In addition, more evidence for HGT is being found in other cases, such as HGT between Bacteria and Eukarya (Watkins and Gary 2006; Guljamow et al. 2007), and within Eukayrya (Nedelcu et al. 2008). Gene duplication is also an important mechanism in molecular evolution. It usually refers to the duplication of regions of DNA that contain at least one gene (Fig. 1c-A). Gene loss is the loss of DNA sequences of genes (Fig. 1c-B). There are many factors that could lead to gene loss, such as unequal crossing over and losses from translocation. Both gene duplication and gene loss are very common and their mechanisms have been extensively studied (Dittmar and Liberles 2011).

**Figure 1:**
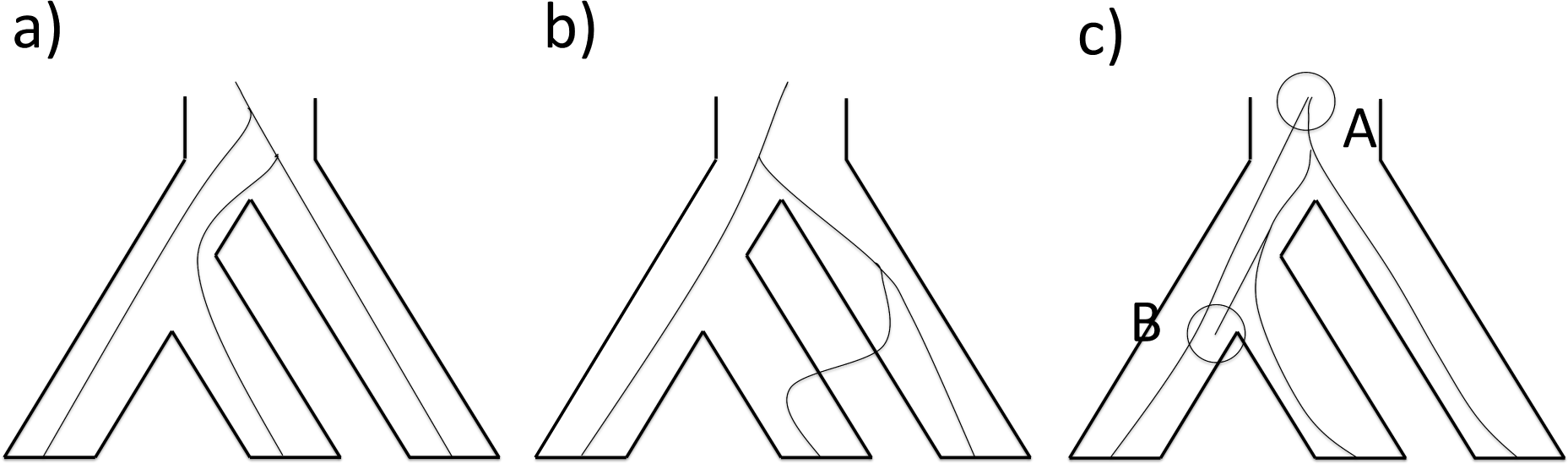
Factors responsible for the incongruence of gene trees and the species tree. a) Deep coalescence. b) Gene flow. c) A-gene duplication, B-gene loss.

## Coalescent Theory and Deep Coalescence

The best-studied of these processes is deep coalescence, mainly because Kingman’s coalescent theory, a continuous-time retrospective model in which the genes sampled in individuals can be traced back to a common ancestor known as the most recent common ancestor (MRCA), provides a way to model the in-population coalescent processes and to link them on a phylogenetic tree. To illustrate this idea, we consider a three-taxon species tree *S* = ((1,2),3) with only one lineage sampled from each population. There are three possible gene tree topologies ((1,2),3), ((2,3),1) and ((1,3),2), with four possible gene histories *H_a_*, *H_b_*, *H_c_*, and *H_d_* (see Figs. 2a∼d). According to Kingman (1982a,b), in a population with *θ* = 4*N μ*, where *N* is the effective population size and *μ* is the mutation rate, the random variable *T*, defined to be the time for *n* lineages to coalesce into *n* − 1 lineages, follows an exponential distribution with the parameter 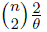. In our case, let *T* be the time to the coalescent event of the two lineages sampled from population 1 and population 2 (see Fig. 2a). Assume 2*/θ*_12_ = 1 so that *T* ∼ *Exp*(1) in the population 12. Let *t* be the time interval between the two population divergence events. Then the probability of the gene history *H_a_* is equal to the probability that *T* is smaller than or equal to *t*. So the probability of gene history *H_a_* can be expressed as a function of *t*,

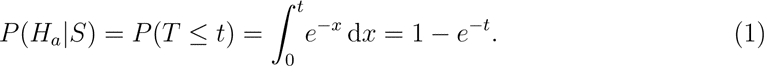

**Figure 2:**
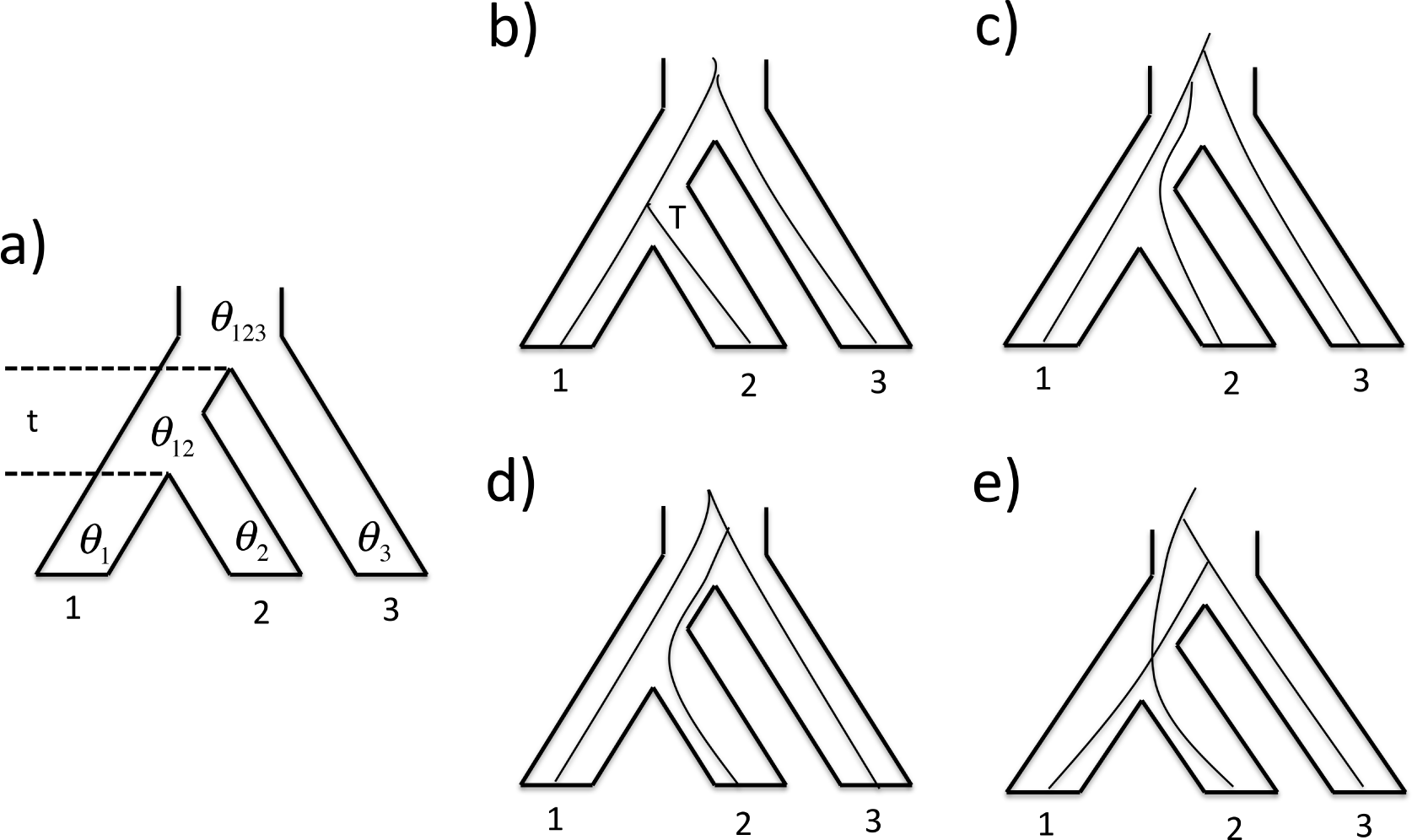
Different gene histories given a three-taxon species tree. a) *G_a_*: linages sampled from population 1 and 2 coalesce in the ancestral population 12 during the time interval t. b) *G_b_*: both coalescent events happen in the ancestral population 123 with gene tree ((1,2),3). c) *G_c_*: both coalescent events happen in the ancestral population 123 with gene tree ((2,3),1). d) *G_d_*: both coalescent events happen in the ancestral population 123 with gene tree ((1,3),2).

Since mating is random in the population 123, *G_b_*, *G_c_* and *G_d_* all have the same probability (Figs. 2b,c,d),

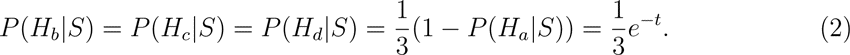

Therefore, the probability of gene tree *G* given species tree *S* is

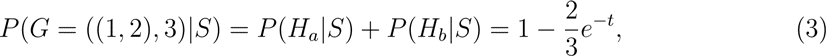

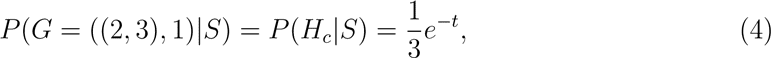

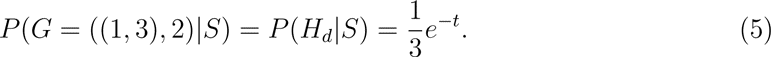

Equations 3, 4, and 5 can be used to calculate the distribution of gene trees *G* given a species tree *S*. On the other hand, species trees can be estimated by examing the gene tree distribution. Based on similar ideas, a number of methods have been proposed to estimate species trees with the assumption that the conflict between gene trees and species trees are due solely to deep coalescence.

*BEAST (Drummond and Rambaut 2007; Drummond et al. 2012) and BEST (Bayesian Estimation of Species Trees Under the Coalescent Model)(Liu 2012) are two widely-used Bayesian inference programs. *BEAST uses Makov Chain Monte Carlo (MCMC) to jointly estimate the posterior distribution of the target species tree as well as all the gene trees and other population parameters such as mutation rates and population sizes. BEST deploys a hierarchical MCMC approach for the same purpose. The program STEM (Species Tree Estimation Using Maximum Likelihood) (Kubatko et al. 2009) takes a set of gene trees as the input and returns the maximum tree (MT) as an estimate of their underlying species tree. Liu (2006; see also Liu and Pearl, 2010) has shown that MT is statistically consistent if the gene trees are known. This method was also developed independently by Mossel and Roch (2010) under the name GLASS (Global LAteSt Split). STEM can be viewed both as a maximum likelihood method and as a distance method, since MT is a maximum likelihood estimate under suitable conditions but it can be built from a distance matrix where each entry is the smallest coalescent time of genes from every pair of species.

## The IM Model and Gene Flow

There are more phylogenetic inference programs that model deep coalescence than those listed here (See Felsenstein’s website http://evolution.genetics.washington.edu/phylip/software.html/ for a more complete list). However, how to appropriately model gene flow for species tree estimation has been less well-studied and still remains a big challenge. Maddison (1997) described the gene tree parsimony method that picks the species tree with the minimal number of migration events, and then a decade later, Eckert and Carstens (2008) and Leache et al. (2014) examined the accuracy of species tree estimates from simulated data subject to gene flow for several of the existing species tree estimation methods, none of which models the process of gene flow. They concluded that the existence of migration may complicate the phylogenetic inference problem in many situations. Kutschera et al. (2014) also confirmed this conclusion in an empirical data study.

The IM (Isolation-with-Migration) model, which can be used to calculate the probability density of coalescent times in the presence of gene flow, may be the key tool to solve the problem. To introduce the IM model, we consider a two-population IM model that involves six parameters, *θ*_1_, *θ*_2_, *θ_A_*, *τ*, *m*_12_ and *m*_21_ (Fig. 3). We define *θ_i_* = 4*N_i_μ*, *i* ∈ {1, 2, *A*} where *N_i_* is the effective population size for the corresponding population *i* and *μ* is the mutation rate per generation. *τ* is the speciation time (the length of the time interval from the time of speciation to the present). We further define *m_ij_* = *M_ij_/μ*, where *M_ij_* is the migration rate from population *i* to population *j* per generation. Assume that one lineage is sampled from each population. Let the state *S*_(*i,j*)_ indicate *i* genes in population 1 and *j* genes in population 2. We can enumerate all the possible states before time *τ* (if not specifically mentioned time is always viewed from present to past throughout the text): *S*_(1,1)_ (also the initial state), *S*_(2,0)_, *S*_(0,2)_, *S*_(1,0)_, *S*_(0,1)_. To formulate an instantaneous rate matrix, the transition rate between every pair of states needs to be calculated. For example, Figure 3b-A illustrates the transition from state *S*_(1,1)_ to state *S*_(2,0)_ through a migration event from population 2 to population 1, which has rate *m*_21_. Figure 3b-B is the transition from state *S*_(2,0)_ to state *S*_(1,0)_ through a coalescent event in population 1, which has a rate of 2*/θ*_1_. Figure 3b-C is the transition from state *S*_(1,0)_ to state *S*_(0,1)_ through a migration event, which has rate *m*_12_. The instantaneous rate matrix **Q** is given below:

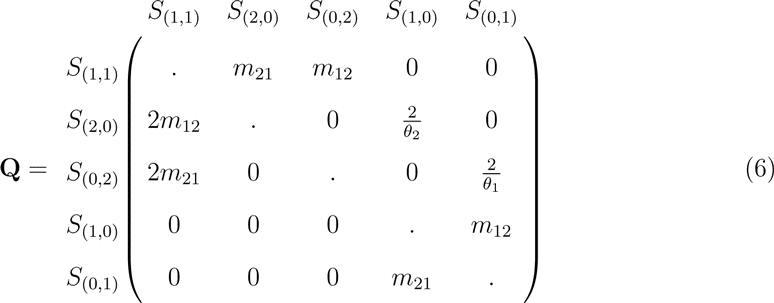

**Figure 3:**
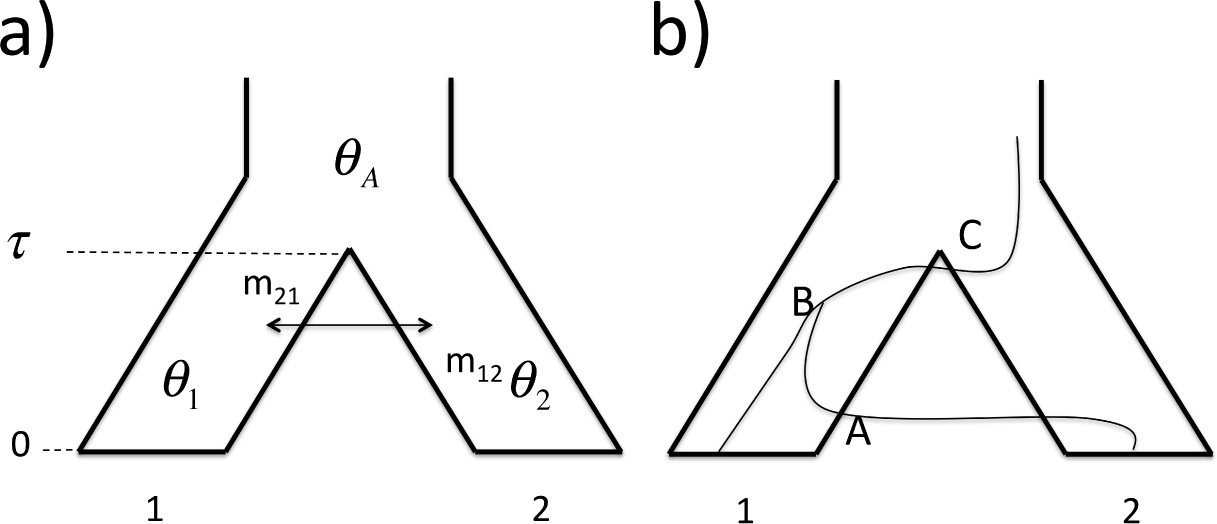
A two population IM model. a) Model species tree and parameters. b) Illustration of state change: A. S(1,1) to S(2,0) by migration, B. S(2,0) to S(1,0) by coalescence, C. S(1,0) to S(0,1) by migration.

The diagonal entries are filled in so that the sum of each row is zero. After time *τ*, there are 2 possible cases:

I. There is only one lineage in the ancestral population, which means the coalescent event has happened before time *τ*. Therefore, the state at time *τ* could be either *S*_(0,)_ or *S*_(1,0)_.
II. There are two lineages at time *τ* in the ancestral population. The state at time *τ* could be *S*_(0,2)_, *S*_(1,1)_ or *S*_(2,0)_.

Following Hobolth et al. (2011), the continuous-time Markov chain representation can be used to get the matrix of probabilities of transitions between the states as a function of time. This transition probability matrix is obtained as the solution **P**(*t*) to the system of differential equations **P′**(*t*) = **QP**(*t*) with initial condition **P**(0) = **I**. The solution is **P**(*t*) = *e***^Q^***^t^*, which we use to derive the probability density function for the two cases listed above. Let t be the time to the coalescent event.

<u>Case I:</u> Note that this case corresponds to *t* ≤ *τ*, and we must consider two possibilities:

1. If the coalescent event occurs in population 1, the density for time t ≤ *τ* is

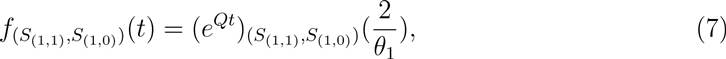

where 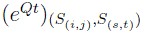 is the entry (*a, b*) in the matrix *e^Qt^* if *S*_(*i,j*)_ is in the *a^th^* row and *S*_(*s,t*)_ is in the *b^th^* column.
2. If the coalescent event occurs in population 2, the density for time t ≤ *τ* is

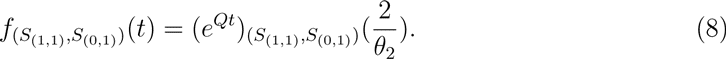

<u>Case II:</u> If the coalescent event occurs after time *τ*, the density for *t* > *τ* is

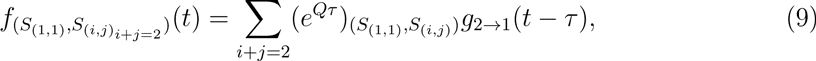

where *g_n→_*_1_(*y*) is the probability density function for *n* genes to coalesce to 1 gene in time *y*. This probability is well-known (Tavaré 1984; Takahata and Nei 1985; Wakeley 2009; Efromovich and Kubatko 2008). The special case 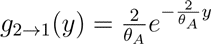 follows from basic coalescent theory.

Equations 7 - 9 can be used to derive formulas to calculate the distribution of gene trees given a species tree with the presence of gene flow in the same way that we have previously. Questions arise whether a similar approach is appliable to estimate species trees in the presence of gene flow. At least so far, such a phylogenetic inference program hasn’t been developed. Nontheless, the IM model has already been widely used in demographic parameter estimation. Hey and Nielsen (2004) developed a Bayesian program called IM under an IM model, which is the first software to jointly estimate speciation times, population sizes and migration rates. The upgraded versions are IMa (2007) and IMa2 (2010). Zhu and Yang (2012) also developed an IM-model-based likelihood method to jointly estimate speciation times, population sizes and migration rates. All of these methods assume that the correct species tree topology is known. In order to study populations with gene flow, many researchers first obtain a species tree estimate by a species tree estimation program that doesn’t allow the possibility of gene flow. Then they treat this species tree estimate as the correct phylogeny and use a demographic parameter inference program to evaluate the magnitude of migration. However, they are risking the chance that errors in the species tree estimation may also collapse the demographic parameter estimation.

In this paper, we propose a distance method called STEST (Species Tree Estimation from Speciation Times) to estimate species trees in the presence of gene flow. The idea is to use pairwise speciation time or species divergence time estimates as distances to construct a species tree. Species tree estimation error is not involved in a two-species case, where there is only one possible species tree topology. Therefore pairwise speciation times are first estimated by a speciation time estimation method that assumes the possbility of gene flow. Then a sequential clustering algorithm is applied to construct the species tree. Despite the fact that the motivation to develop this method is to accomodate gene flow, we also evaluate the performance of our method using a speciation time estimation method that assumes no gene flow.

## Methods

Our method STEST consists of two parts: creation of a distance matrix and use of a clustering algorithm based on this matrix to construct the tree. We will describe each of these steps in detail in the following sections.

### Speciation Time Estimation Methods

To create a distance matrix, a speciation time estimation method needs to be picked first. Here, we prefer maximum likelihood methods over Bayesian methods for their short computation time. Three different likelihood methods are considered: Yang (2002)’s method M0, Yang (2010)’s method M1, and Zhu and Yang (2012)’s method SIM3s. M1 and SIM3s allow the possbility of gene flow while M0 does not. All three estimate the parameters of interest by searching a point in the parameter space that maximizes the likelihood function. The following is a brief description of the formulation of their likelihood functions.

#### SIM3s

We start with the method SIM3s because the other two methods can be illustrated under the SIM3s model’s setting (see Fig. 4a): there are three populations 1, 2 and 3 (outgroup); *τ*_0_ and *τ*_1_ are speciation times; *θ_i_* = 4*N_k_μ* (*k* = 1, 2, 3, 12, 123) are the population size parameters with *N*’s being the effective population sizes and *μ* the mutation rate per site; gene flow is assumed to exist between population 1 and population 2 with migration rates *m*_12_ and *m*_21_; assume that *θ*_1_ = *θ*_2_ = *θ* and *m*_12_ = *m*_21_= *m*, and that one lineage is sampled from each species. By convention, Θ = (*θ, θ*_12_*, θ*_123_*, m, τ*_0_*, τ*_1_) is the parameter vector. Under this setting, there are 6 possible gene histories, *H*_1*a*_, *H*_1*b*_, *H*_1*c*_, *H*_1*d*_, *H*_2_ and *H*_3_ (Figs. 4b∼g). That gene flow only exists between population 1 and population 2 before time *τ*_1_ fits a two-population IM model. By setting *m*_12_ = *m*_21_ = *m*, *θ*_1_ = *θ*_2_ = *θ*, *τ* = *τ*_1_ and *θ_A_* = *θ*_12_, formulas 6 - 9 in the introduction can be used. Let *t*_1_, *t*_0_ be the times to the first and to the second coalescent events, respectively. We can derive the probability density *f* (*t*_1_*, t*_0_*, H|*Θ) of coalescent times *t*_1_, *t*_0_ and gene history *H* given Θ (for convenience, we write *f* (*t*_1_*, t*_0_*, H*) instead in the cases where no ambiguity arises).

**Figure 4:**
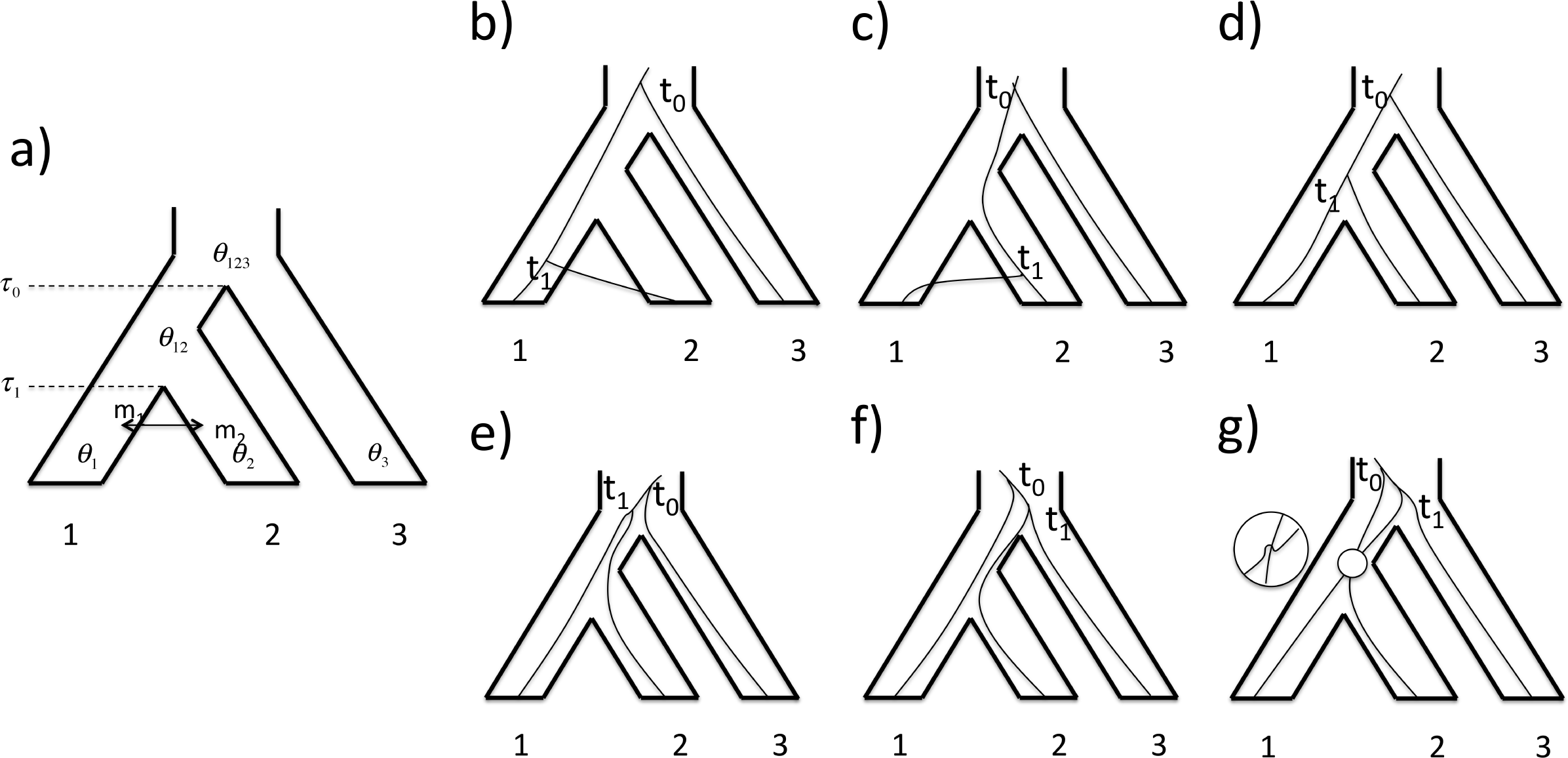
Zhu and Yang’s SIM3s model. a) Model species tree and parameters. b) *H*_1*a*_: Coalescence of 1,2 happens first in population 1, *t*_1_ *≤ τ*_1_. c) *H*_1*b*_: Coalescence of 1,2 happens first in population 2, *t*_1_ *≤ τ*_1_. d) *H*_1*c*_: Coalescence of 1,2 happens first in population 12, *τ*_1_ *≤ t*_1_ *≤ τ*_0_. e) *H*_1*d*_: Coalescence of 1,2 happens first in population 123, *τ*_0_ *≤ t*_1_ *≤ t*_0_. f) *H*_2_: Coalescence of 2,3 happens first in population 123, *τ*_0_ *≤ t*_1_ *≤ t*_0_. g) *H*_3_: Coalescence of 1,3 happens first in population 123, *τ*_0_ *≤ t*_1_ *≤ t*_0_.

For gene history *H*_1*a*_ (Fig. 4b), *t*_1_< *τ*_1_, the first coalescent event occurs in population 1 and the second coalescent event occurs in population 123, so

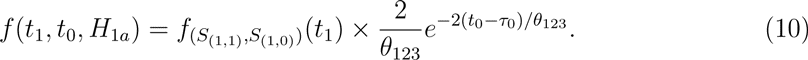

For gene history *H*_1*b*_ (Fig. 4c), *t*_1_ < *τ*_1_, the first coalescent event occurs in population 2 and the second coalescent event occurs in population 123, so

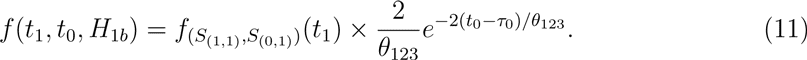

For gene history *H*_1*c*_ (Fig. 4d), *τ*_1_ < *t*_1_ < *τ*_0_, the first coalescent event occurs in population 12 and the second coalescent event occurs in population 123, so

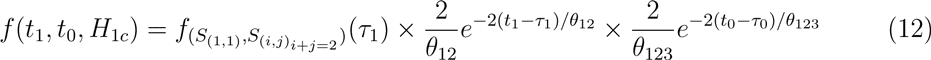

For gene history *H*_1*d*_, *H*_2_ and *H*_3_ (Figs. 4e,f,g), *t*_1_ > *τ*_0_ and both coalescent events occurs in population 123, so

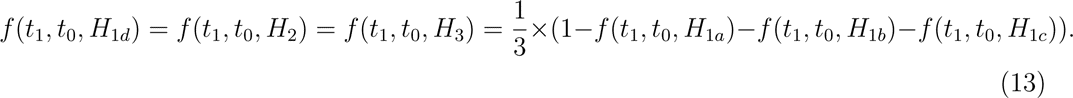

Suppose the sequences at each locus are aligned and no gaps exist. At each site, there are five possible patterns: *xxx*, *xxy*, *yxx*, *xyx* and *xyz*, where *x*, *y*, *z* are symbols for different nucleotides. At any locus *i*, the sequence alignments are first summarized into site pattern counts 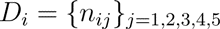, where *n_ij_* is the number of the j^th^ site pattern observed. Let 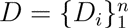 be the data for *n* unlinked loci and assume the JC69 mutation model (Jukes and Cantor 1969). Equations 10 - 13 and Yang (1994)’s formula for the conditional probability *P* (*D_i_|t*_1_*, t*_0_*, H*) of *D_i_* given *t*_1_, *t*_2_ and *H* are then combined together to derive the likelihood function of Θ given *D*,

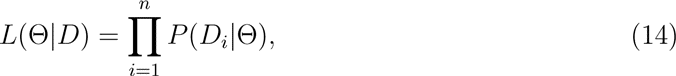

where

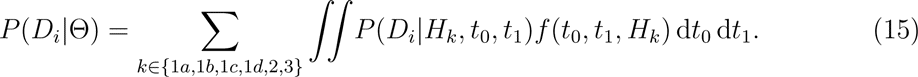

#### M0

The method M0 adopts a reduced model of SIM3s, which assumes no gene flow. Under this setting, there are only 4 possible gene histories: *H*_1*c*_, *H*_1*d*_, *H*_2_ and *H*_3_ (Fig. 4). This is also the case for the coalescent without migration (see Fig. 2). The likelihood function is the same as Equation 14 with *m* ≡ 0. Let *f*_0_(*t*_1_*, t*_0_*, H*) denote the probability density of *t*_1_, *t*_0_ and *H* in this special case when *m* ≡ 0. Then by Equations 10- 13,

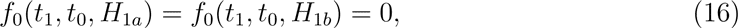

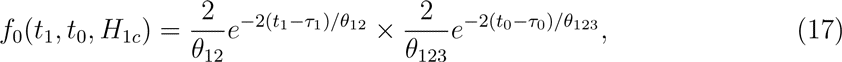

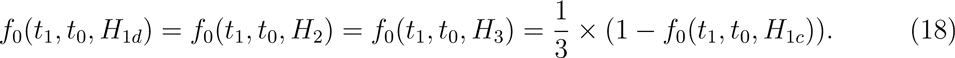

The likelihood function therefore can be written as

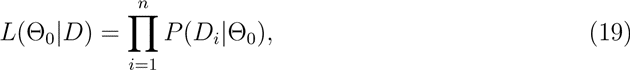

where

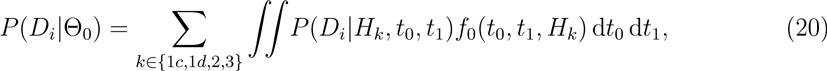

and where the parameter vector Θ_0_ = (*θ*_12_*, θ*_123_*, τ*_1_*, τ*_0_) because *θ* does not affect the likelihood value, thus is unidentifiable.

#### M1

The difference between M1 and M0 is that M1 allows the species divergence time *τ*_1_ of species 1 and species 2 to vary among loci at random due to possible gene flow. Yang (2010) chooses a beta distribution to model this. The density of *τ*_1_ is

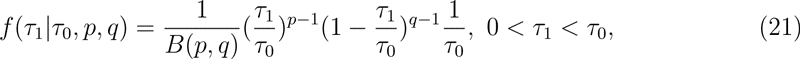

where *τ*_0_, *p*, and *q* are prameters of the distribution. He then changes variables by making 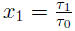. Then *x*_1_ ∼ *beta*(*p, q*), 0 < *x*_1_ < 1. The mean 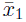 of *x*_1_ is 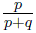 and the variance is 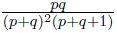, so 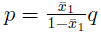. Treating 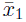 and *q* as the parameters of the distribution of *x*_1_, the density of *x*_1_ can be written as 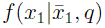. The likelihood function is

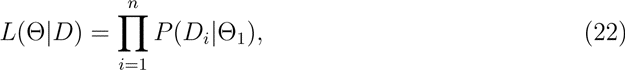

where

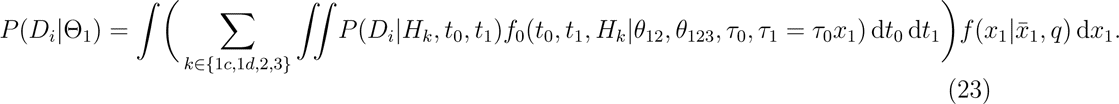

and where the parameter vector 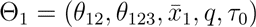. The estimate of the speciation time *τ*_1_ is 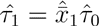.

We denote our method STEST (SIM3s) if SIM3s is used to estimate speciation times. STEST (M0) and STEST (M1) are defined similarly. Once a speciation time estimation method is picked, the distance matrix can be built easily.

### Distance Matrix Building

Let 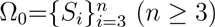 be a set of species. Let *S*_0_ be the outgroup. Within each species *S_i_*(0 ≤ *i* ≤ *n*), multiple genes *<u>gi</u>* are sampled. For each pair of species (*S_i_*, *S_j_*), 0 < *i* ≠ *j*, (<u>*g_i_*</u>, <u>*g_j_*</u>, <u>*g*_0_</u>) are used to estimate the speciation time *t_i,j_* between species *S_i_* and *S_j_* using one of the methods M0, M1 or SIM3s. We define *D* = (*t_i,j_*) to be the distance matrix, which is a symmetric *n* × *n* matrix that contains the speciation times for all pairs of species.

### Species Tree Reconstruction

Let *T*_0_ = {*t_i,j_*}*_i<j_* be the set of all of the entries of the lower triangular part of the distance matrix D, i.e., distance between every pair of species. The following algorithm is performed:

1. Pick the smallest time 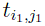 in *T*_0_, write 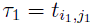, add a new node at time *τ*_1_ to connect 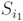 and 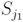.
2. Suppose *k* nodes have been added and a set Ω ⊂ Ω_0_ of species has been connected on the tree. Pick the smallest time 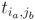 among the remaining times.

Case 1. If 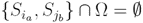, add a new node at time 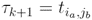 connecting 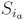 and *S_jb_*.
Case 2. If 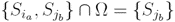, then add a new node at time 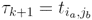 connecting 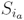 and the node at 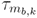, where 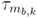 is the largest time at which the node is connected to *S_jb_* after *k* nodes have been added. Similarly for the case in which 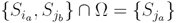.
Case 3. If 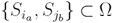, then (*i*) if 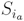 and 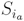 share an ancestor, then discard the time 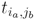, this step is finished; (*ii*) if 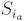 and 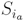 don’t share an ancestor, add a new node at time 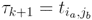 to connect the nodes at 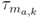 and 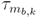.
3. Continue until all species share a common ancestor, i.e. the root is reached.

### An Example

To illustrate this method, we consider a set *S* = {*S*_1_*, S*_2_*, S*_3_*, S*_4_*, S*_5_} consisting of 5 species. Let *D* = (*t_i,j_*) be the distance matrix, for example,

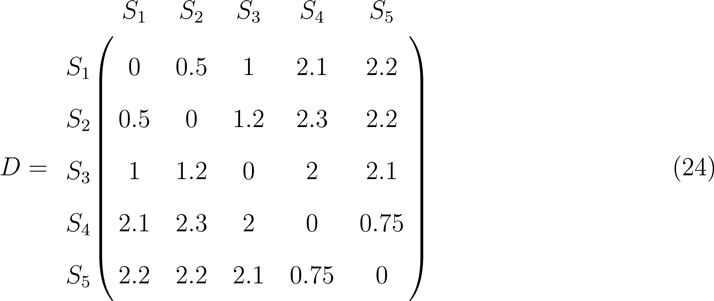

Then *T*_0_ = {*t_i,j_*}*_i<j_* = {0.5, 0.75, 1, 1.2, 2, 2.1, 2.2, 2.3}. We perform the clustering algorithm step by step (see Fig. 5).

1. Pick the samllest element in *T*_0_, *t*_1,2_ = 0.5. Add a new node at time *τ*_1_ = *t*_1,2_ = 0.5 to connect *S*_1_ and *S*_2_. After this step, Ω = {*S*_1_*, S*_2_}, *T* = *T*_0_–{0.5}.
2. Pick the smallest element in *T*, *t*_4,5_ = 0.75. Since 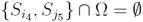, add a new node at time *τ*_2_ = *t*_4,5_ to connect *S*_4_ and *S*_5_. After this step, Ω = {*S*_1_*, S*_2_*, S*_4_*, S*_5_}, *T* = *T*_0_–{0.5, 0.75}.
3. Pick the smallest element in *T*, *t*_1,3_ = 1, then {*S*_1_*, S*_3_} ∩ Ω = {*S*_1_}, and *τ_m_*_1,2_ = *τ*_1_. Add a new node at time *τ*_3_ to connect *S*_3_ and the node at *τ*_1_. After this step, Ω = {*S*_1_*, S*_2_*, S*_3_*, S*_4_*, S*_5_}, *T* = *T*_0_–{0.5, 0.75, 1}.
4. Pick the smallest element in *T*, *t*_2,3_ = 1, but {*S*_2_*, S*_3_} ⊂ Ω and *S*_2_ & *S*_3_ share a common ancestor at *τ*_2_ so nothing is done. After this step, Ω = {*S*_1_*, S*_2_*, S*_3_*, S*_4_*, S*_5_}, *T* = *T*_0_–{0.5, 0.75, 1, 1.2}.
5. Pick the smallest element in *T*, *t*_3,4_ = 2. Since {*S*_3_*, S*_4_} ⊂ Ω and *S*_3_ and *S*_4_ do not share a common ancestor, we need to add a new node to connect the nodes at *τ_m_*_3,3_ and *τ_m_*_4,3_, i.e., nodes at *τ*_3_ and *τ*_2_. After this step, Ω = {*S*_1_*, S*_2_*, S*_3_*, S*_4_*, S*_5_}, *T* = *T*_0_–{0.5, 0.75, 1, 1.2, 2}.
6. Root is reached!

**Figure 5:**
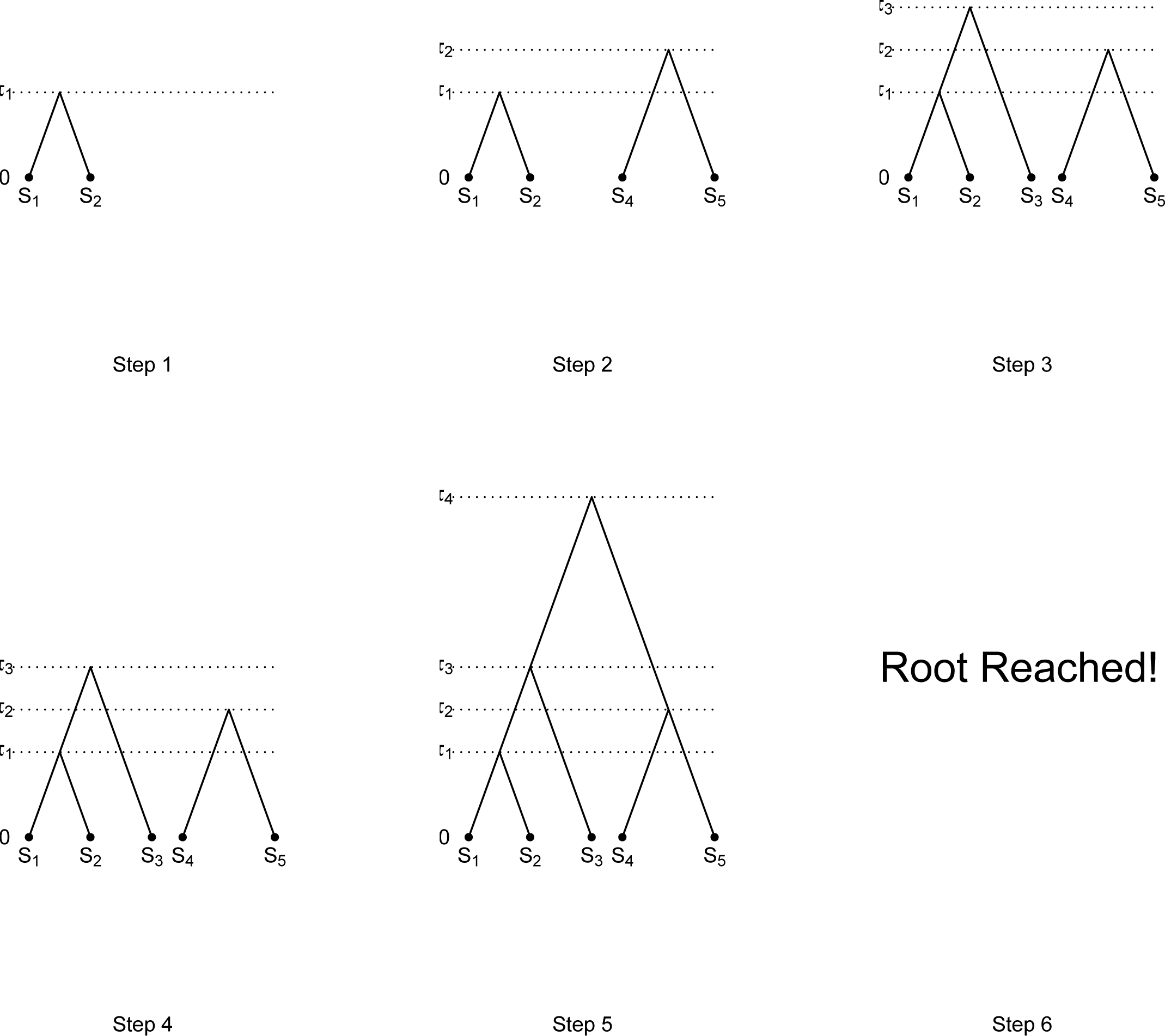
Illustration of tree reconstruction algorithm.

This algorithm can be easily implemented in R. We analyze both simulated data and empirical data to evaluate the performance of our methods.

### Simulation Study

#### Simulation Study 1: Four-taxon Tree Under the n-island Model

Figure 6 shows the model species tree and parameters for the first simulation study. 100 genes are sampled with one lineage sampled from each species under a four-taxon species tree. Since the methods M0, M1 and SIM3s all require an outgroup, we specify species 0 to be the outgroup. All of the population size parameters are assumed to be the same and equal to *θ* = 4. Three bifurcating speciation events happen at times *τ*_1_, *τ*_2_, and *τ*_3_. Gene flow exists among all but population 0 before *τ*_1_ with *m_ij_* being the migration rate from population *i* to population *j* (1 ≤ *i, j* ≤ 3). All the migration rates are equal. Thus the migration pattern follows an n-island model (Wright 1943).

**Figure 6:**
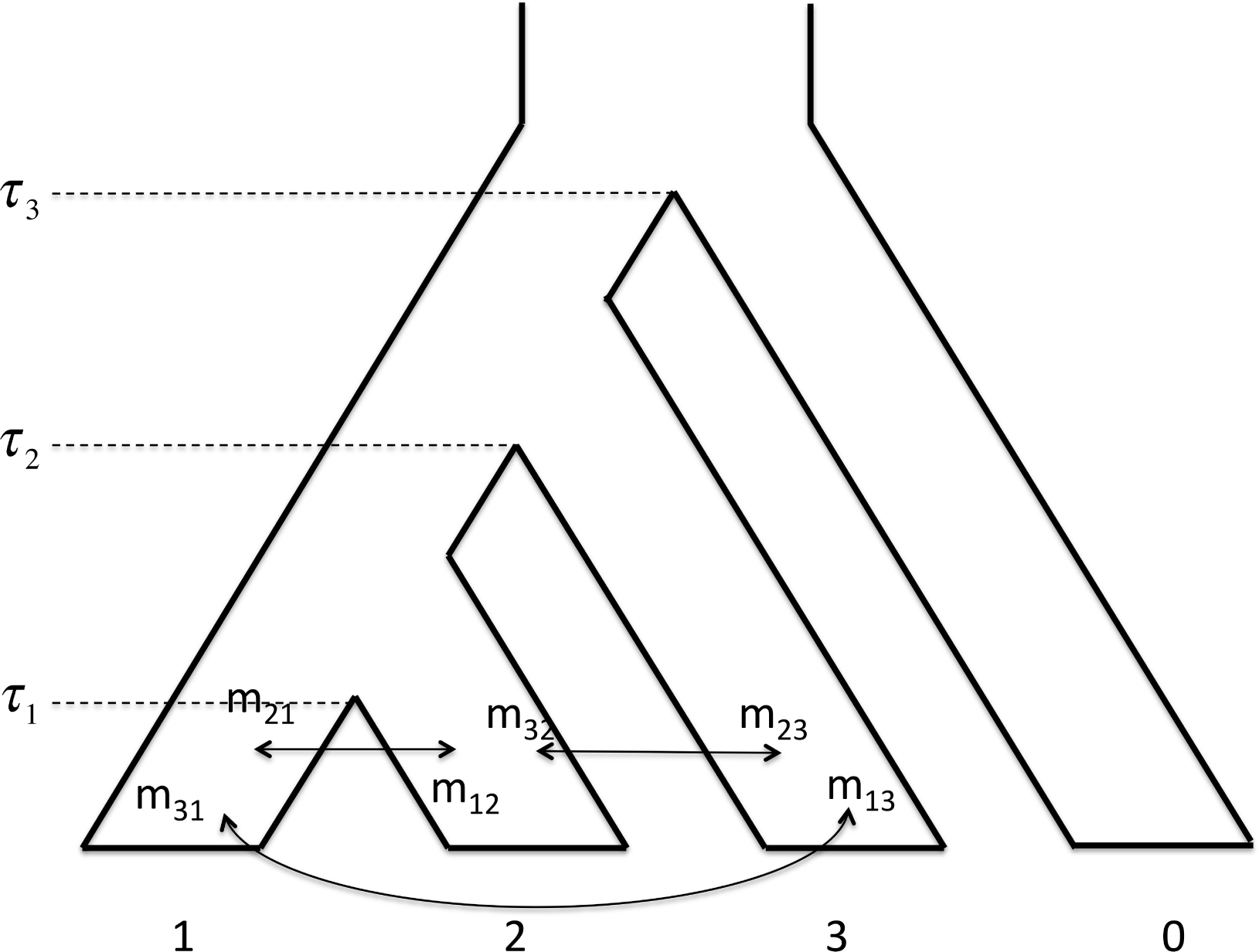
Model species tree and parameters for simulation study 1.

To simulate the data, gene trees are first sampled from ms (Hudson 2002) under 18 different settings (labeled by A1∼A9 and B1∼B9, see Table 1). Seq-Gen (Rambaut and Grassly 1997) is then used to generate full sequence data from the simulated gene trees under the JC69 model (Jukes and Cantor 1969). The length of each gene is set to be 1, 000 bp. STEST (M0), STEST (M1), STEST (SIM3s) and *BEAST (for 8 settings as indicated in Table 1) are used to get species tree estimates directly from sequence data. Gene trees are first estimated by PAUP* (Swofford 2002) using maximum likelihood (ML) and are then used as the input to STEM. For each setting, the same procedure is repeated 80 times.

**Table 1:**
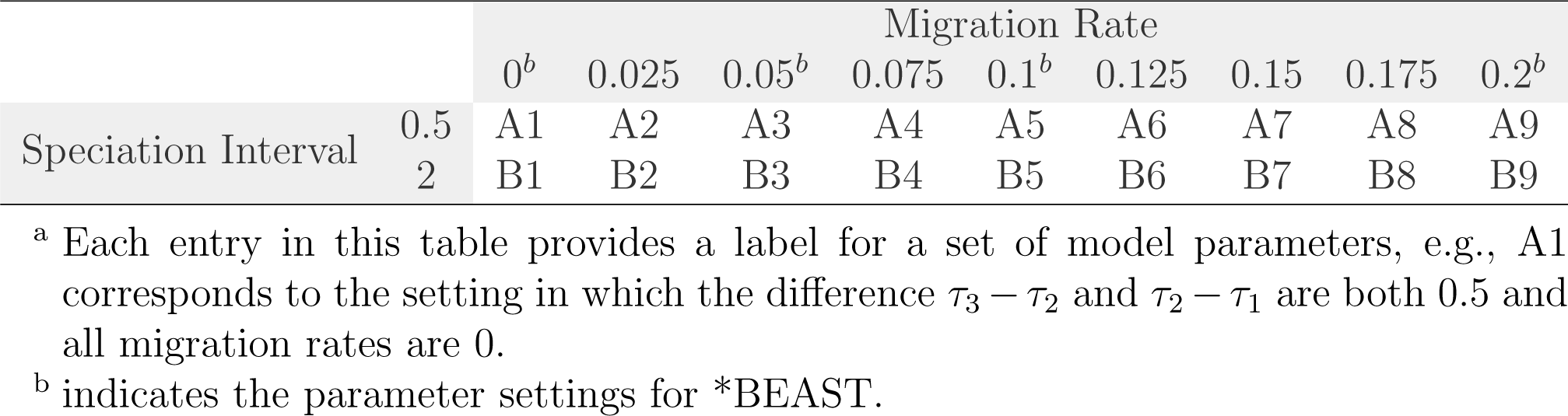
Settings for simulation study 1*^a^*.

#### Simulation Study 2: Nine-taxon Tree Under the n-island Model

Figure 7 shows the model species tree and parameters for the second simulation study. 100 genes are sampled with one lineage sampled from each species under a nine-taxon species tree. We specify species 0 to be the outgroup. All of the population size parameters are assumed to be the same and equal to *θ* = 4. Eight bifurcating speciation events happen at times *τ*_1_, *τ*_2_, *τ*_3_, *τ*_4_, *τ*_5_, and *τ*_6_. Gene flow exists among all but population 0 before *τ*_1_ with *m_ij_* the migration rate from population *i* to population *j* (1 *≤ i, j ≤* 3). All of the migration rates are assumed to be equal. Thus the migration pattern again follows an n-island model.

**Figure 7:**
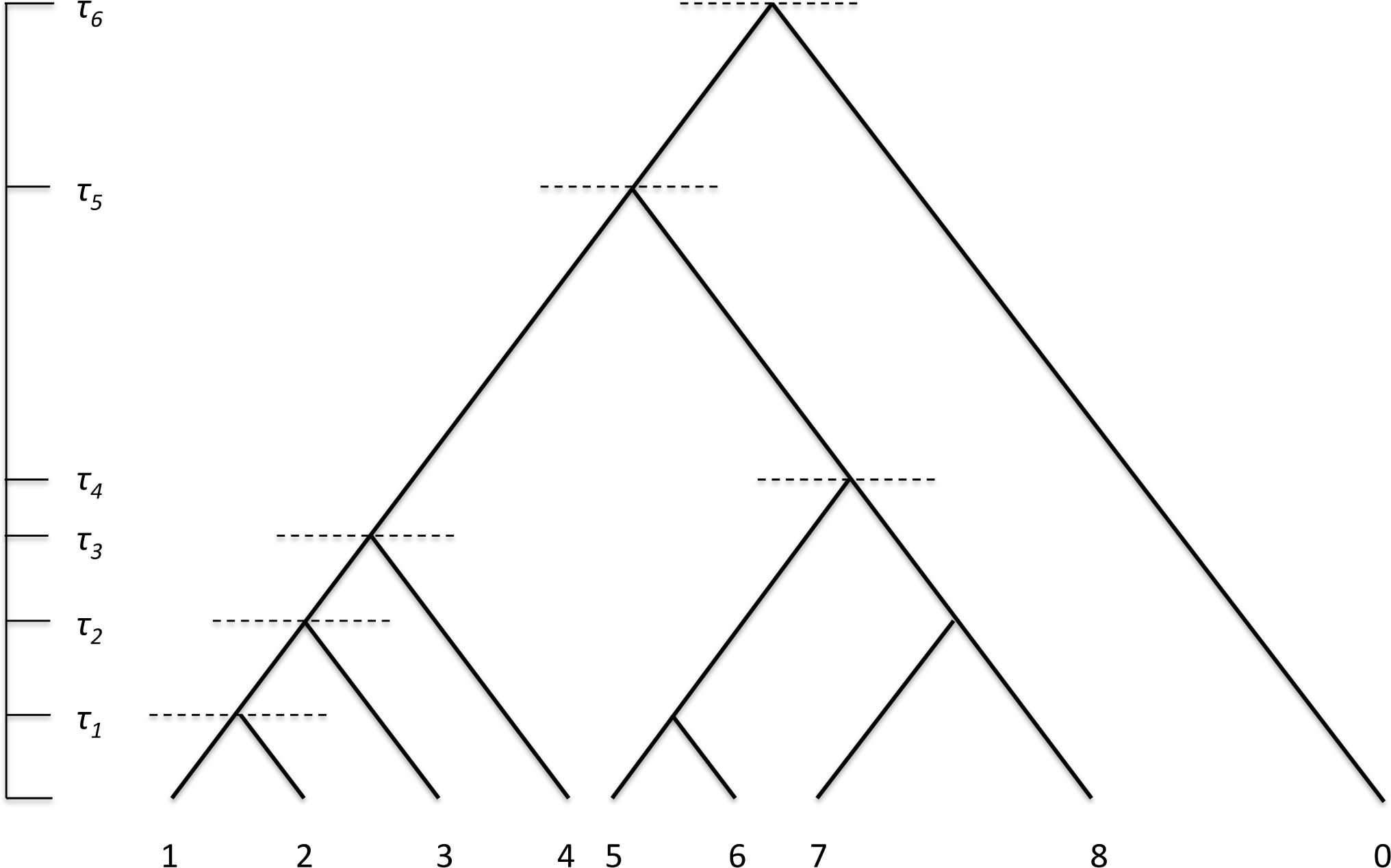
Model species tree and parameters for simulation study 2.

Just as in the previous section, gene trees are sampled from ms and then Seq-Gen is used to generate sequence data from these simulated gene trees under the JC69 model. The length of the simulated sequences is 1, 000 bp. STEST (M0), STEST (M1) and STEST (SIM3s) are applied to the sequence data directly. PAUP* is used to estimate ML gene trees from sequence data before STEM is applied. The 18 parameter settings C1∼C9 and D1∼D9 are listed in Table 2. We use 100 replicates for each setting.

**Table 2:** Settings for simulation study 2*^a^*.

|  | Migration Rate |  |  |  |  |  |  |  |  |
| --- | --- | --- | --- | --- | --- | --- | --- | --- | --- |
|  | 0 | 0.025 | 0.05 | 0.075 | 0.1 | 0.125 | 0.15 | 0.175 | 0.2 |
| $\tau_1 = 1, \tau_2 = 2, \tau_3 = 3, \tau_4 = 4, \tau_5 = 5, \tau_6 = 10$ | C1 | C2 | C3 | C4 | C5 | C6 | C7 | C8 | C9 |
| $\tau_1 = 2, \tau_2 = 3, \tau_3 = 4, \tau_4 = 5, \tau_5 = 6, \tau_6 = 10$ | D1 | D2 | D3 | D4 | D5 | D6 | D7 | D8 | D9 |

<sup>a</sup> Each entry in this table provides a label for a set of model parameters, e.g., C1 corresponds to the setting in which the speciation times are $\tau_1 = 1, \tau_2 = 2, \tau_3 = 3, \tau_4 = 4, \tau_5 = 5, \tau_6 = 10$ and all of the migration rates are 0.

### Empirical Study

We apply the methods STEST (SIM3s), STEST (M0) and STEST (M1) to the HGCOR (Human, Chimpanzee, Gorilla, Orangutan and Rhesus) data set obtained by Ebersberger et al. (2007), who have shown that the species tree topology is ((((G,O),C),H),R). R (Rhesus) is the outgroup. The data set contains 28, 160 sequence alignments. 249 of sequence alignments are longer than 1, 000 bp and are used in this analysis. This data set is of interest for analysis with these methods, since Zhu and Yang (2012) recently reported gene flow following speciation among some of these taxa.

## Results

### Results for Simulation Study 1

Results from the simulation study 1 are given in Tables 3 and 4 and are plotted in Figure 8. For each setting, the number of correct tree estimates (out of 80) is recorded and translated into a percentage.

**Figure 8:**
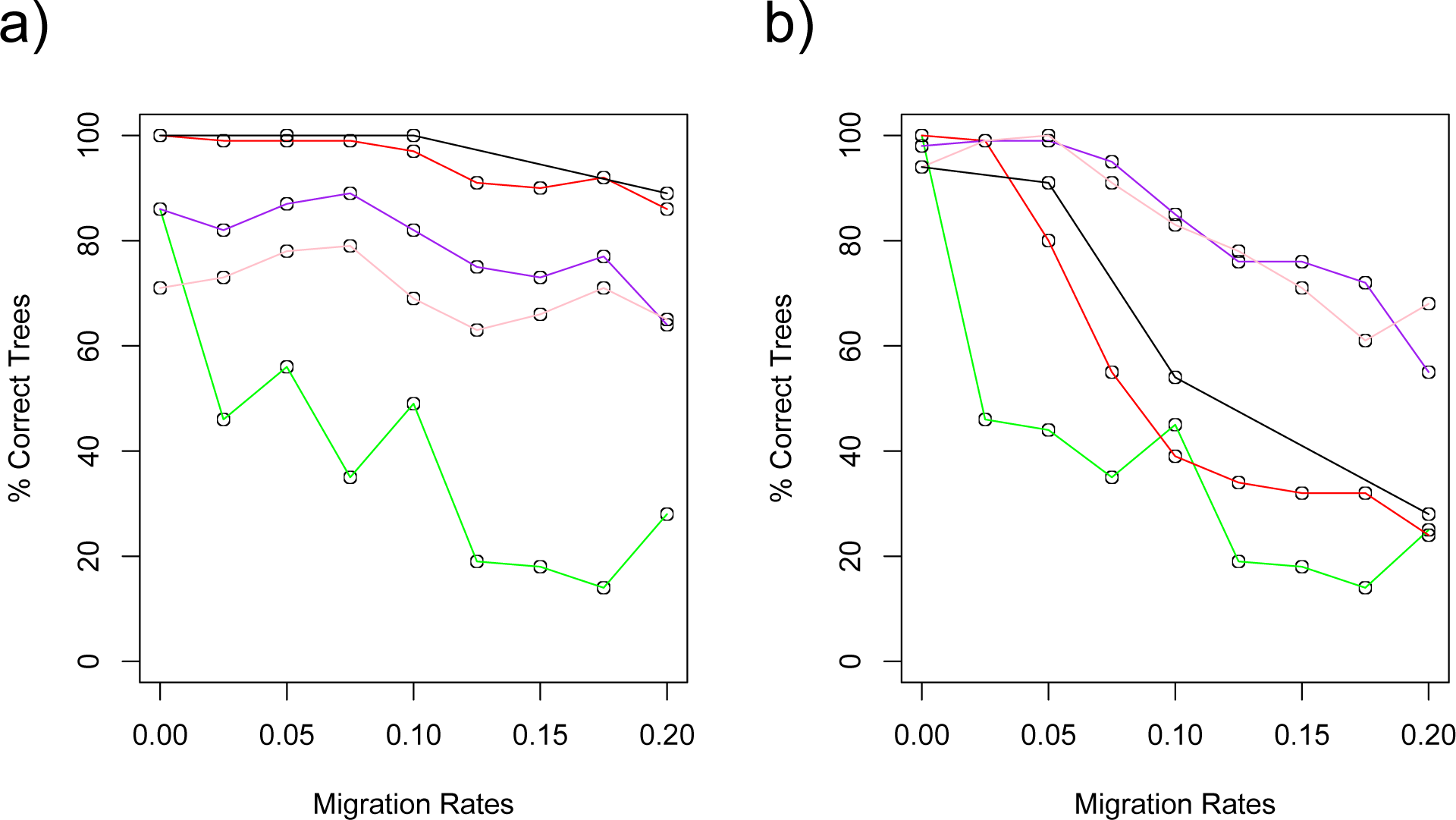
Results plot for simulation study 1. Black: Results from *BEAST; Red: Results from STEST (M0); Purple: Results from STEST (M1); Pink: Results from STEST (SIM3s); Green: Results from STEM. a) is the percentage of the correct estimates vs. the magnitude of the gene flow used to generate data in the short speciation interval case (A1∼A9). b) is the percentage of the correct estimates vs. the magnitude of the gene flow used to generate data in the long speciation interval case (B1∼B9).

**Table 3:** Simulation 1 results (short speciation interval scenario).

| %<sup>a</sup> | A1 | A2 | A3 | A4 | A5 | A6 | A7 | A8 | A9 |
| --- | --- | --- | --- | --- | --- | --- | --- | --- | --- |
| *BEAST | 100 |  | 100 |  | 100 |  |  |  | 89 |
| STEM | 85 | 46 | 56 | 35 | 49 | 19 | 18 | 14 | 28 |
| STEST (M0) | 100 | 99 | 99 | 99 | 97 | 91 | 90 | 92 | 86 |
| STEST (M1) | 86 | 83 | 87 | 89 | 82 | 75 | 73 | 77 | 64 |
| STEST (SIM3s) | 71 | 73 | 78 | 79 | 69 | 63 | 66 | 71 | 65 |

<sup>a</sup> Each entry is the percentage of the correct estimates.

**Table 4:** Simulation 1 results (long speciation interval scenario).

| $\%^a$ | B1 | B2 | B3 | B4 | B5 | B6 | B7 | B8 | B9 |
| --- | --- | --- | --- | --- | --- | --- | --- | --- | --- |
| *BEAST | 94 |  | 91 |  | 54 |  |  |  | 28 |
| STEM | 100 | 46 | 44 | 35 | 45 | 19 | 18 | 14 | 25 |
| STEST (M0) | 100 | 99 | 80 | 55 | 39 | 34 | 32 | 32 | 24 |
| STEST (M1) | 98 | 99 | 99 | 95 | 85 | 76 | 76 | 72 | 55 |
| STEST (SIM3s) | 94 | 99 | 100 | 91 | 83 | 78 | 71 | 61 | 68 |

<sup>a</sup> Each entry is the percentage of the correct estimates.

#### A1∼A9

In the short speciation interval scenarios, *BEAST and STEST (M0) have very similar performance (percentage of correct estimates *>* 85%), which is better than all of the other methods in all cases. When gene flow doesn’t exists, STEM estimates 85% of the total trees correctly, which is close to STEST (M1)’s 86% correct, and better than STEST (SIM3s)’s 71% correct. In the presence of gene flow, the STEST methods consistently yield better results than STEM. The performance of all of the methods decreases as the migration rate increases. Particularly, STEM’s estimation accuracy drops more dramtically than all of the other methods (decreases in accuracy from 85% to 46% when the migration rate changes from 0 to 0.025). The accuracy curves of *BEAST and STEST (M0) are almost flat and remain in a high level (percentage of correct estimates *≥* 97%) when the migration rate is not larger than 0.10. Then they start to drop slowly as the migration rate increases but are still above 85% when the migration rate is increased to 0.20. The performance of STEST (M1) and STEST (SIM3s) follows a similar trend. STEST (M1) performs better than STEST (SIM3s) with a ∼10% difference in accuracy and STEST (M0) is better than STEST (M1) with a ∼15% difference.

#### B1∼B9

In the long speciation interval scenarios, all the methods peform well when there is no gene flow (percentage of correct estimates *≥* 94%), and start to perform worse as the migration rates increases. Again, STEM’s performance decreases dramatically as the migration rate increases (decreases in accuracy from 100% to 46% when the migration rate changes from 0 to 0.025). Similar thing happens to *BEAST and STEST (M0) with the steepest drop in their performance occuring when the migration rate changes from 0.05 to 0.10. STEST (M0) performs better than *BEAST when the migration rate is smaller than 0.05 and the opposite happens when the migration rate is larger than 0.05. STEST (M1) and STEST (SIM3s) outperform all the other methods under every setting. Their performance curves behave similarly to each other, which are almost flat and remian in a high level (percentage of correct estimates *>* 90%) until the migration rate is increased to Their accuracy is still above 50% even when the migration rate is increased to 0.20.

In both scenarios, STEM always performs the worst in the presence of gene flow. In the short speciation interval senarios, *BEAST and STEST (M0) outperforms the other methods and their performance curves are very close to each other. In the long speciation interval scenarios, the same thing happens to STEST (M1) and STEST (SIM3s). The performance curves in the long speciation interval scenarios are steeper than in the short speciation interval scenarios.

### Results for Simulation Study 2

Results from the simulation study 2 are given in Tables 5 and 6 and are plotted in Figures 9 - S8.

**Table 5:** Simulation 2 results.

|  |  | Settings |  |  |  |  |  |  |  |  |
| --- | --- | --- | --- | --- | --- | --- | --- | --- | --- | --- |
|  |  | C1 | C2 | C3 | C4 | C5 | C6 | C7 | C8 | C9 |
| Number of<br>Correct Estimates<br>(Out of 100) | STEM | 87 | 0 | 0 | 1 | 0 | 0 | 0 | 0 | 0 |
|  | STEST (M0) | 100 | 97 | 89 | 78 | 57 | 40 | 29 | 22 | 9 |
|  | STEST (M1) | 93 | 92 | 83 | 80 | 75 | 61 | 68 | 62 | 47 |
|  | STEST (SIM3s) | 86 | 81 | 76 | 68 | 56 | 45 | 43 | 35 | 26 |
| Average RF Distances<br>Between All Estimates<br>and the Correct Tree | STEM | 0.18 | 6.96 | 9.24 | 9.64 | 10.01 | 10.61 | 10.64 | 10.28 | 10.60 |
|  | STEST (M0) | 0 | 0 | 0 | 0.10 | 0.23 | 0.75 | 1.07 | 2.32 | 2.62 |
|  | STEST (M1) | 0.12 | 0.14 | 0.37 | 0.36 | 0.46 | 0.70 | 0.57 | 0.76 | 0.83 |
|  | STEST (SIM3s) | 0.32 | 0.41 | 0.50 | 0.74 | 1.03 | 1.48 | 1.52 | 1.95 | 2.17 |
| Average RF Distances<br>Between Incorrect Estimates<br>and the Correct Tree<sup>a</sup> | STEM | 2.00 | 7.25 | 9.24 | 9.84 | 10.01 | 10.61 | 10.64 | 10.28 | 10.6 |
|  | STEST (M0) |  |  |  | 3.33 | 2.56 | 3.95 | 3.82 | 5.16 | 4.37 |
|  | STEST (M1) | 2.00 | 2.00 | 2.18 | 2.12 | 2.19 | 2.12 | 2.59 | 2.24 | 2.31 |
|  | STEST (SIM3s) | 2.29 | 2.28 | 2.17 | 2.31 | 2.34 | 2.69 | 2.76 | 3.00 | 3.06 |

<sup>a</sup> Missing value means not applicable.

**Table 6:** Simulation 2 results.

|  |  | Settings |  |  |  |  |  |  |  |  |
| --- | --- | --- | --- | --- | --- | --- | --- | --- | --- | --- |
|  |  | D1 | D2 | D3 | D4 | D5 | D6 | D7 | D8 | D9 |
| Number of<br>Correct Estimates<br>(Out of 100) | STEM | 78 | 0 | 0 | 0 | 0 | 0 | 0 | 0 | 0 |
|  | STEST (M0) | 100 | 75 | 56 | 22 | 9 | 1 | 0 | 0 | 0 |
|  | STEST (M1) | 96 | 94 | 84 | 71 | 60 | 45 | 33 | 23 | 18 |
|  | STEST (SIM3s) | 75 | 74 | 69 | 69 | 60 | 45 | 29 | 25 | 11 |
| Average RF Distances<br>Between Estimates and<br>the Correct Tree | STEM | 0.26 | 10.49 | 11.09 | 11.15 | 11.24 | 11.39 | 11.10 | 10.94 | 11.36 |
|  | STEST (M0) | 0 | 0.03 | 0.37 | 2.02 | 3.67 | 6.40 | 7.89 | 9.07 | 9.10 |
|  | STEST (M1) | 0.04 | 0.06 | 0.15 | 0.22 | 0.33 | 0.81 | 1.50 | 1.69 | 2.42 |
|  | STEST (SIM3s) | 0 | 0.02 | 0.14 | 0.33 | 0.47 | 1.05 | 1.80 | 2.04 | 3.09 |
| Average RF Distances<br>Between Incorrect Estimates<br>and the Correct Tree<sup>a</sup> | STEM | 2.00 | 10.49 | 11.09 | 11.15 | 11.24 | 11.39 | 11.10 | 10.94 | 11.36 |
|  | STEST (M0) |  | 3.00 | 3.70 | 4.30 | 5.17 | 6.88 | 8.05 | 9.16 | 9.10 |
|  | STEST (M1) | 2.00 | 3.00 | 2.50 | 2.20 | 2.06 | 2.70 | 3.00 | 3.02 | 3.78 |
|  | STEST (SIM3s) |  | 2.00 | 2.33 | 2.20 | 2.47 | 2.44 | 2.90 | 3.24 | 3.77 |

<sup>a</sup> Missing value means not applicable.

#### C1∼C9

In the cases when *τ*_1_ = 1, all methods perform well in the presence of gene flow in terms of the percentage of correct estimates (*≥* 86%, see Fig. 9). All methods’ performance decreases as the migration rate increases. STEM’s estimation accuracy quickly goes down to 0% correct when the migration rate is 0.025 while all the other methods still remain above 55% when the migration rate is 0.10. STEST (M0) outperforms all of the other methods when the migration rate is small (0 ∼ 0.075). Its accuracy curve starts to drop below STEST (M1)’s when the migration rate is larger than 0.1 and starts to drop below STEST (SIM3s)’s when the migration rate is larger than 0.125. STEST (M1)’s accuracy is consistently better than STEST (SIM3s) with a ∼15% difference. When the migration rate is increased to 0.20, STEST (M1) performs the best with a 47% accuracy. STEST (SIM3s) is the second best with a 26% accuracy (Table 5 and Fig. 9a).

**Figure 9:**
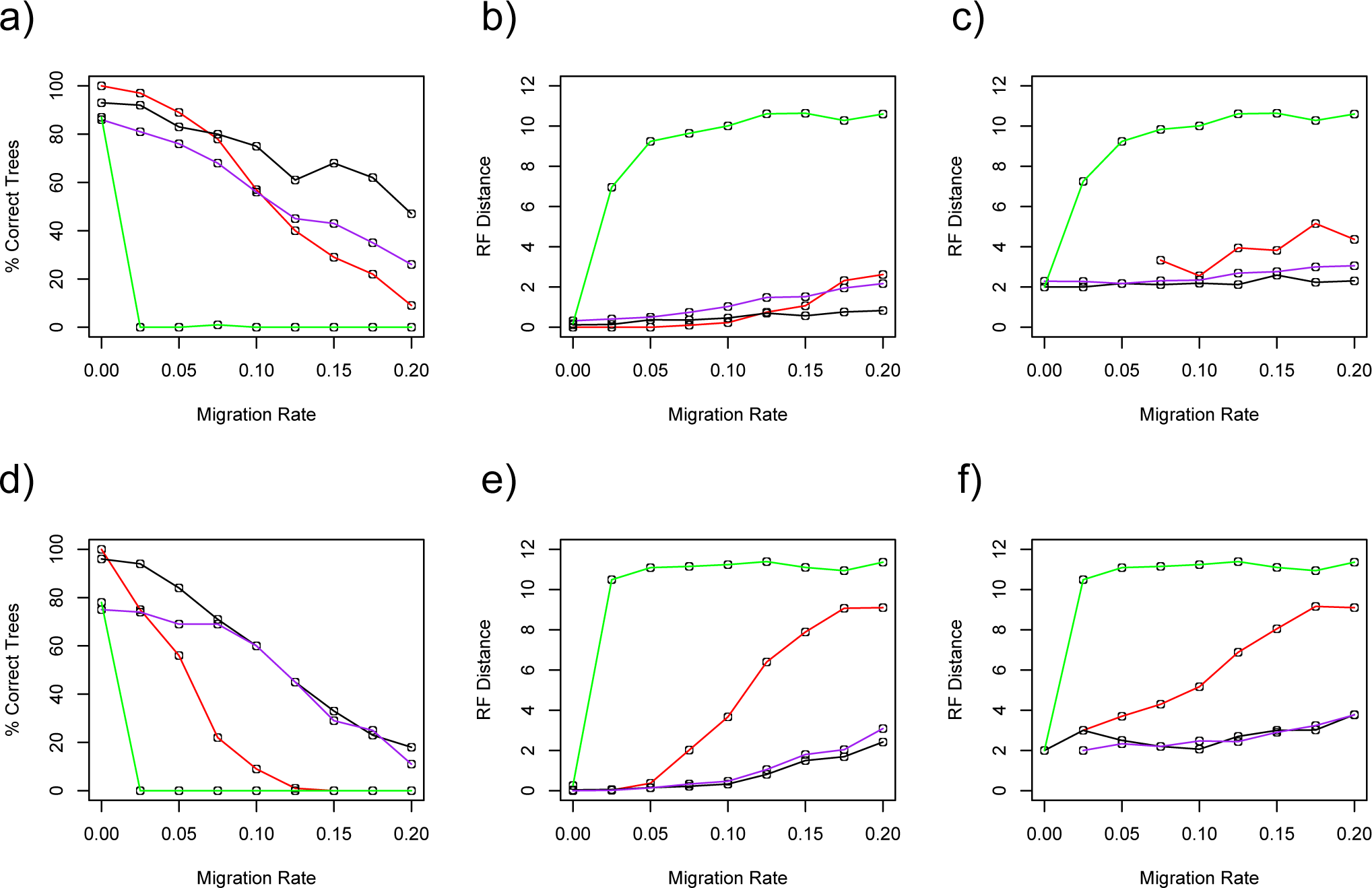
Results plot for simulation study 2. Green: Results from STEM; Red: Results from STEST (M0); Black: Results from STEST (M1); Purple: Results from STEST (SIM3s). a) is the percentage of the correct estimates vs. the magnitude of the gene flow used to generate data for *τ*_1_ = 1 (C1∼C9). b) is the average Robinson-Foulds distances between all estimates and the correct tree vs. the magnitude of the gene flow used to generate data for *τ*_1_ = 1 (C1∼C9). c) is the average Robinson-Foulds distances between incorrect estimates and the correct tree vs. the magnitude of the gene flow used to generate data for *τ*_1_ = 1 (C1∼C9). d) is the percentage of the correct estimates vs. the magnitude of the gene flow used to generate data for *τ*_1_ = 2 (D1∼D9). e) is the average Robinson-Foulds distances between all estimates and the correct tree vs. the magnitude of the gene flow used to generate data for *τ*_1_ = 2 (D1∼D9). f) is the average Robinson-Foulds distances between incorrect estimates and the correct tree vs. the magnitude of the gene flow used to generate data for *τ*_1_ = 2 (D1∼D9).

The **a**verage **R**obinson-**F**oulds **d**istances (RF distances, see Robinson and Foulds, 1981) between **a**ll the estimates and the **c**orrect tree (aRFdac) for different methods and different migration rates are plotted in Figure 9b. Note that RF distances is designed to measure the distances between unrooted trees. It is possible that two different rooted trees have RF distance 0. Nontheless, RF distance is still the most popular metric for rooted trees. All methods’ aRFdac increases as the migration rate increases. Even though STEM’s estimation accuracy decreases to 0 when the migration rate is 0.025, its aRFdac is just 6.98. This distance continues to increase as the migration rate increases. It attains values above 10 when the migration rate is larger than 0.1. STEST (M0)’s aRFdac is the smallest when the migration rate is small (*≤* 0.10). It becomes larger than STEST (M1)’s when the migration rate is larger than 0.125 (0.75 *>* 0.70 when *m* = 0.125), and becomes larger than STEST (SIM3s)’s when the migration rate is larger than 0.175 (2.32 *>* 1.95 when *m* = 1.75). STEST (M1)’s aRFdac is smaller than STEST (SIM3s)’s under all settings. It never exceeds 0.83 and STEST (SIM3s)’s aRFdac never exceeds 2.17 (Table 5).

The **a**verage **RF d**istances between the **i**ncorrect estimates and the **c**orrect tree (aRFdic) for different migration rates are plotted in Figure 9c. All methods’ aRFdic increases as the migration rate increases. STEM’s aRFdic curve is very similar to its aRFdac curve except that the minimum possible value it attains is 2 instead of 0. STEST (M1) and STEST (SIM3s)’s aRFdic increases very slowly as the migration rate increases. Their aRFdic curves are similar to each other. STEST (M1)’s aRFdic never exceeds 2.59 and STEST (SIM3s)’s aRFdic never exceeds 3.06 when migration rate falls in the interval (0, 0.20). STEST (M0)’s aRFdics, ranging from 2.56 to 5.16, is always larger than STEST (M1)’s and STEST (SIM3s)’s.

Figures S1∼S4 are the histograms showing the frequency of the RF distances for species tree estimates using different methods. When migration rate is 0, most of the estimates have zero RF distances to the correct tree. The histogram shows a unimodal distribution with the peak at the RF distance 0 (See Figs. S1a,S2a,S3a,S4a). As the migration rate increases, more and more estimates have large distances to the correct tree. The distribution first becomes multimodal with multiple short peaks or uniform (e.g., see Fig. S1b), and then becomes unimodal again with the peak at a high RF distance value (e.g., see Fig. S1i). This process is observed in STEM shown in Figure S1. All the methods seem to follow a similar trend. For STEST (M0) and STEST (SIM3s), their distributions are about to become multimodal when the migration rate is increased to 0.20 (see Fig. S2i and Fig. S4i). But STEST (M1)’s distribution remains unimodal with the peak at RF distance 0 in all cases we investigate (Fig. S3).

**Figure S1:**
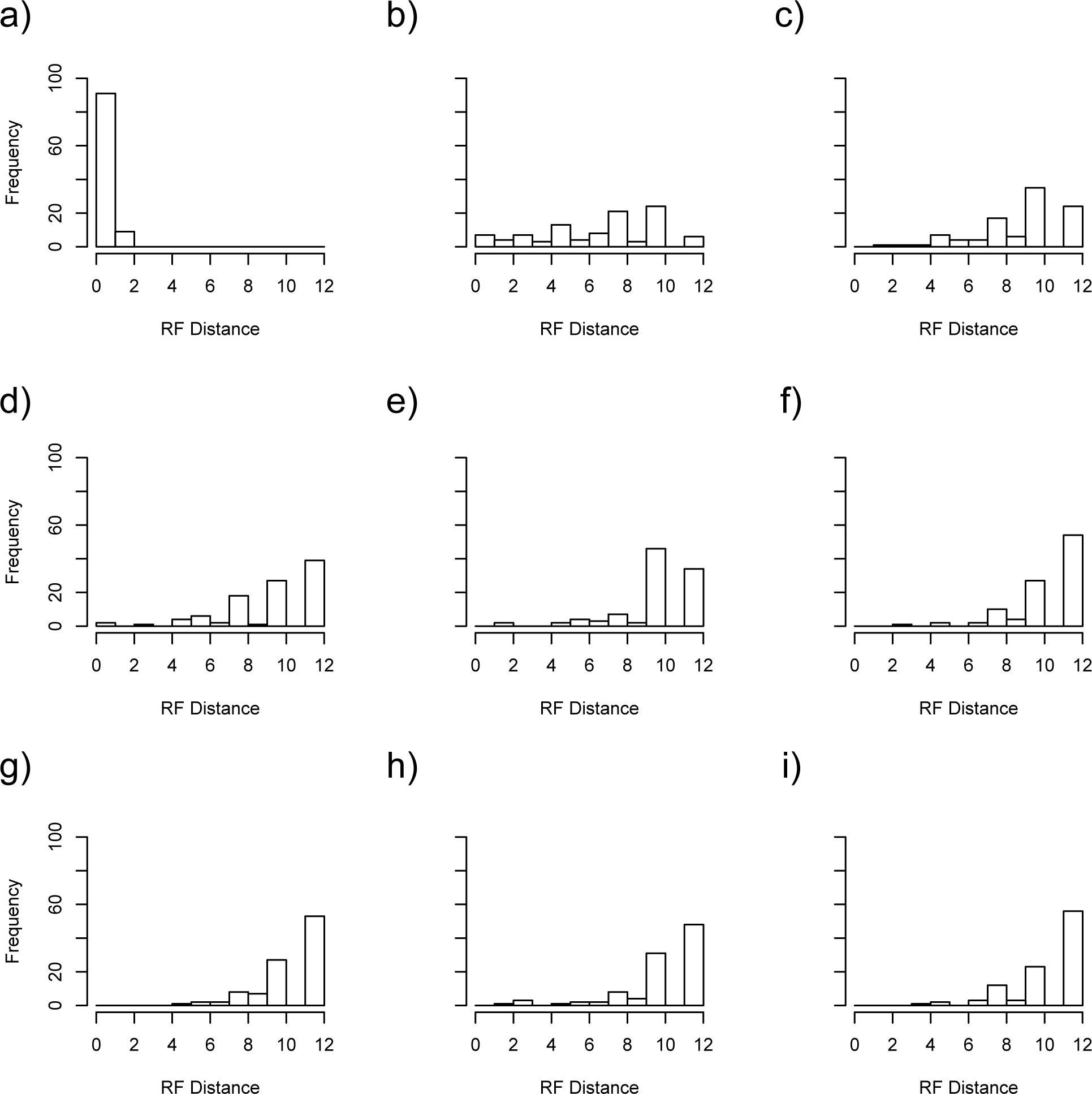
Frequency histogram showing the distribution of Robinson-Foulds distances between estimates using STEM and the correct tree under settings: a) C1, b) C2, c) C3, d) C4, e) C5, f) C6, g) C7, h) C8, and i) C9.

**Figure S2:**
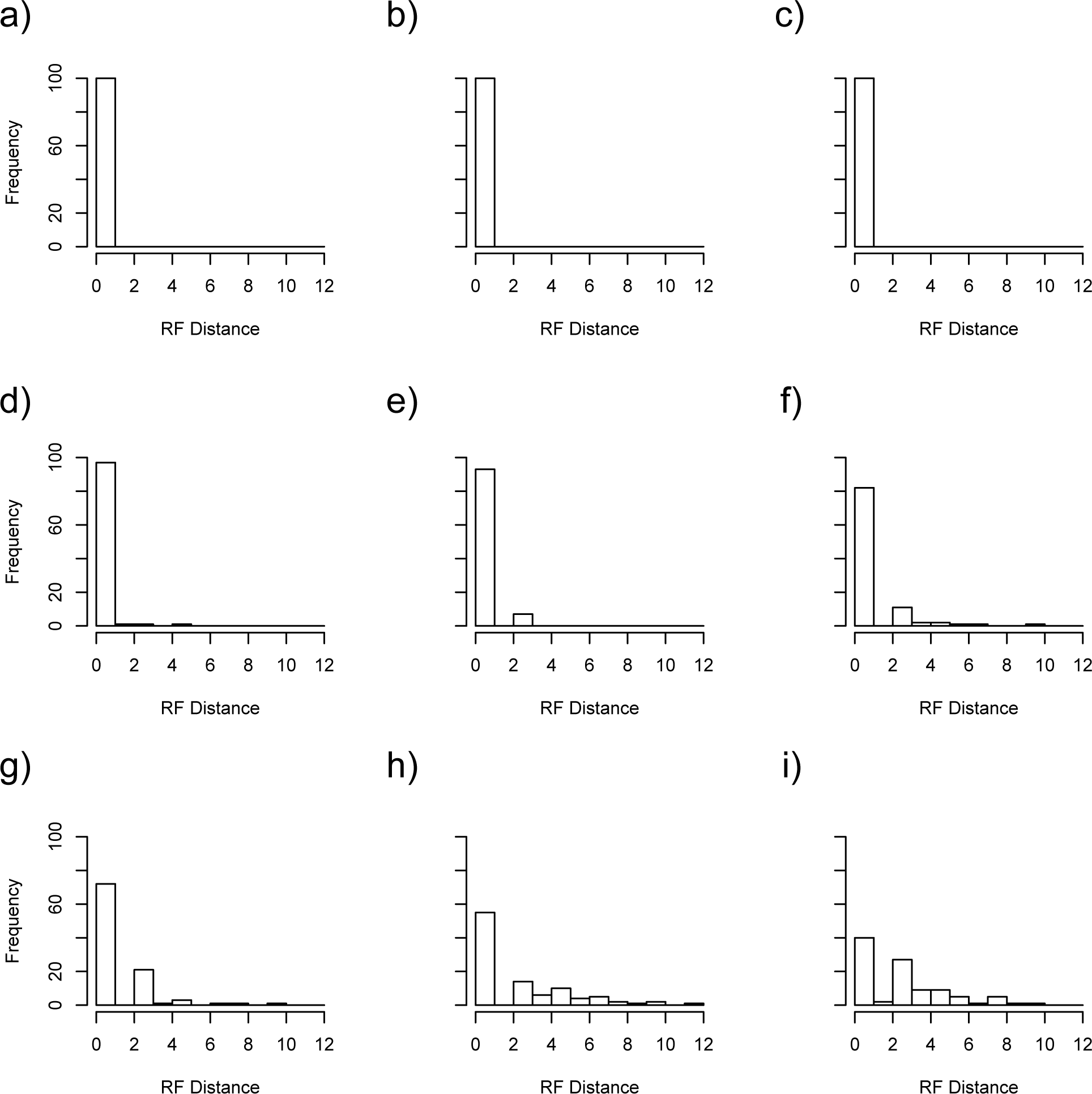
Frequency histogram showing the distribution of Robinson-Foulds distances between estimates using STEST (M0) and the correct tree under settings: a) C1, b) C2, c) C3, d) C4, e) C5, f) C6, g) C7, h) C8, and i) C9.

**Figure S3:**
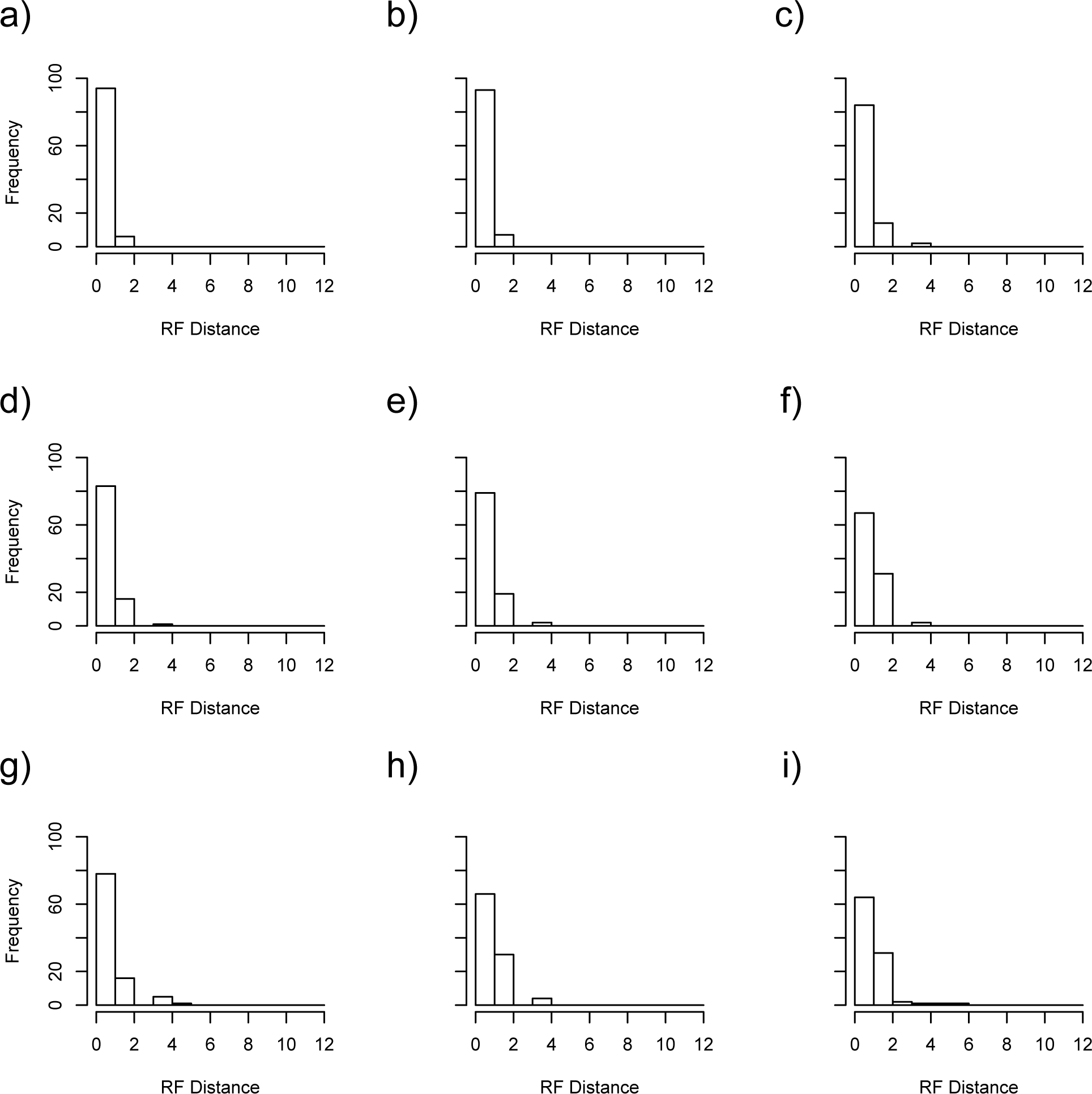
Frequency histogram showing the distribution of Robinson-Foulds distances between estimates using STEST (M1) and the correct tree under settings: a) C1, b) C2, c) C3, d) C4, e) C5, f) C6, g) C7, h) C8, and i) C9.

**Figure S4:**
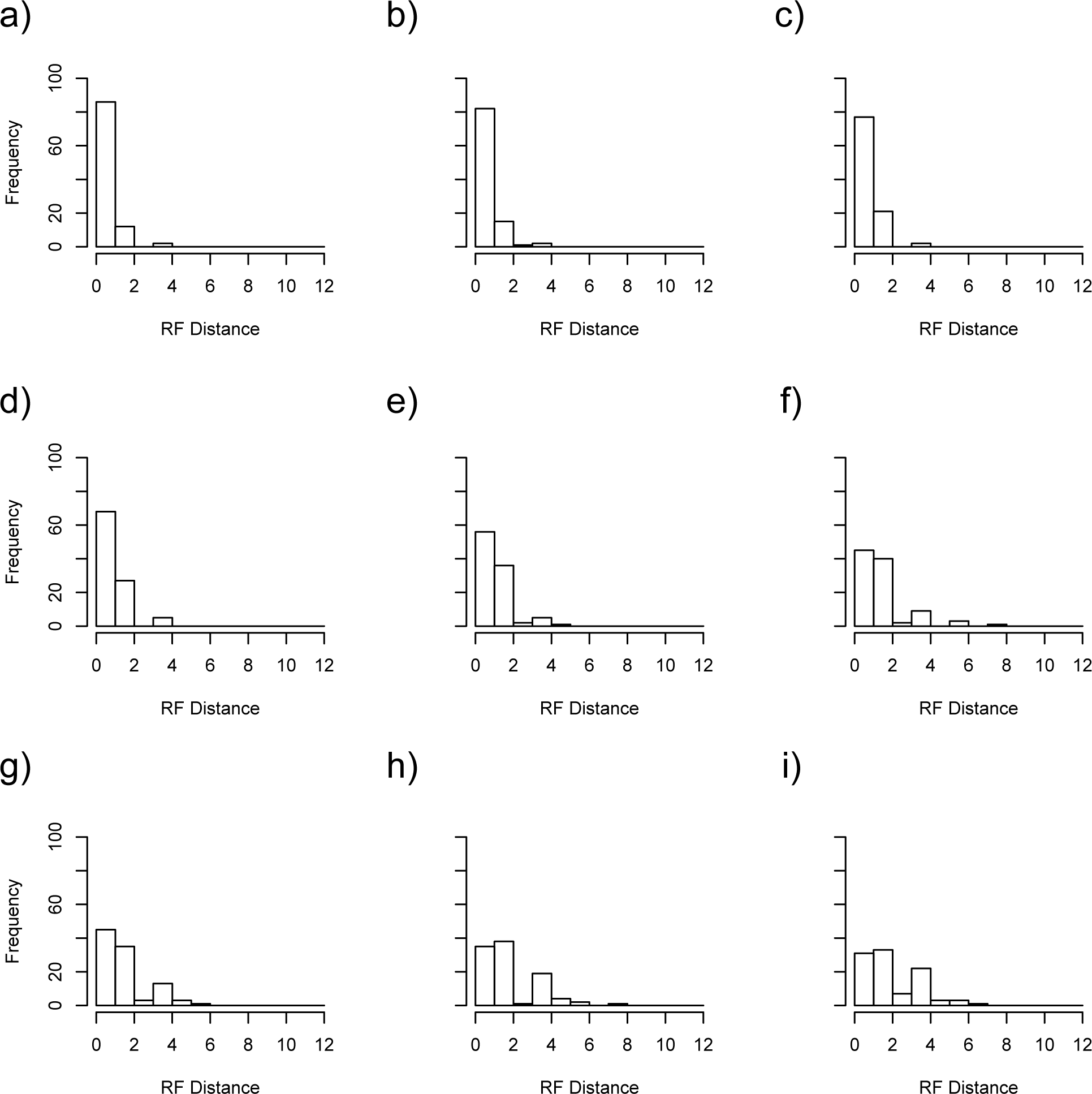
Frequency histogram showing the distribution of Robinson-Foulds distances between estimates using STEST (SIM3s) and the correct tree under settings: a) C1, b) C2, c) C3, d) C4, e) C5, f) C6, g) C7, h) C8, and i) C9.

#### D1∼D9

In the cases when *τ*_1_ = 2, STEST (M0) and STEST (M1) perform very well in the presence of gene flow in terms of the percentage of correct estimates (*≥* 96%, see Fig. 9). STEM and STEST (SIM3s) also performs well with 78% correct and 75% correct, respecitively, in estimation accuracy. All methods’ performance decreases as the migration rate increases. Again, STEM’s estimation accuracy quickly goes down to 0% correct when the migration rate is 0.025. STEST (M1)’s estimation accuracy also decreases dramatically as the migration rate increases. It goes down to 1% at the migration rate 0.125. STEST (SIM3s)’s estimation accuracy stays 69 ∼ 75% when the migration rate is smaller than 0.1. STEST (M1)’s estimation accuracy decreases from 96% to 71% when the migration rate is increased from 0 to 0.075. However, STEST (SIM3s) and STEST (M1)’s performance curves become similar to each other when the migration rate is larger than 0.1. They drop from 60% to below 20% as the migration rate increases from 0.10 to 0.20 (Table 6 and Fig. 9d).

The aRFdac for different methods under different migration rates are plotted in Figure 9e. All methods’ aRFdac increases as the migration rate increases. The trend is more obvious than the previous cases. STEM’s aRFdac attains values above 10 at migration rate as small as 0.025. When the migration rate is smaller than 0.05, STEST (M0), STEST (M1) and STEST (SIM3s)’s aRFdacs are small (*<* 0.4) and do not increase a lot. STEST (M0)’s aRFdac curve starts to have a higher increasing rate when the migration rate increases from 0.05 to 0.20. Its aRFdac value is above 9 when the migration rate is increased to 0.175. STEST (M1) and STEST (SIM3s)’s aRFdac curves are again very similar to each other and increase slowly as the migration rate increases. Their aRFdac values do not exceed 3.09 even when the migration rate is 0.20.

The aRFdic for different methods under different migration rates are plotted in Figure 9f. All methods’ aRFdic increases as the migration rate increases. Again, STEM’s aRFdic curve is very similar to its aRFdac curve except that the minimum possible value it attains is 2 instead of 0. It attains values above 10 at the migration rate 0.025. STEST (M0)’s aRFdac also increases very fast. Its aRFdic value becomes larger than 8 at migration rate 0.15 and larger than 9 at migration rate 0.175. STEST (M1) and STEST (SIM3s)’s aRFdic curves are again similar to each other. Their values increases slowly as the migration rate increases and stay smaller than 3 at migration rate smaller than 0.15 and smaller than 4 at migration rate smaller than 0.20.

Figures S5 – S8 are the histograms showing the frequency of the RF distances for species tree estimates using different methods. Similarly to the cases when *τ*_1_ = 1, most of the estimates have zero RF distances to the correct tree at migration rate 0. The histograms show a unimodal distribution with the peak at the RF distance 0 (See Figs. S5a,S6a,S7a,S8a). As the migration rate increases, more and more estimates have large distances to the correct tree. The distribution first becomes multimodal with multiple short peaks or even uniform (e.g., see Fig. S6f), and then becomes unimodal again with the peak at a high RF distance value (e.g., see Fig. S6i). This whole process is observed in STEST (M0) shown in Figure S6. All methods seem to follow this trend. STEM skips the multimodal or uniform stage. STEST (SIM3s)’s distribution is about to become multimodal when the migration rate is increased to 0.20. The distribution for STEST (M1) remains unimodal with the peak at RF distance 0 under all settings we investigate (Fig. S3).

**Figure S5:**
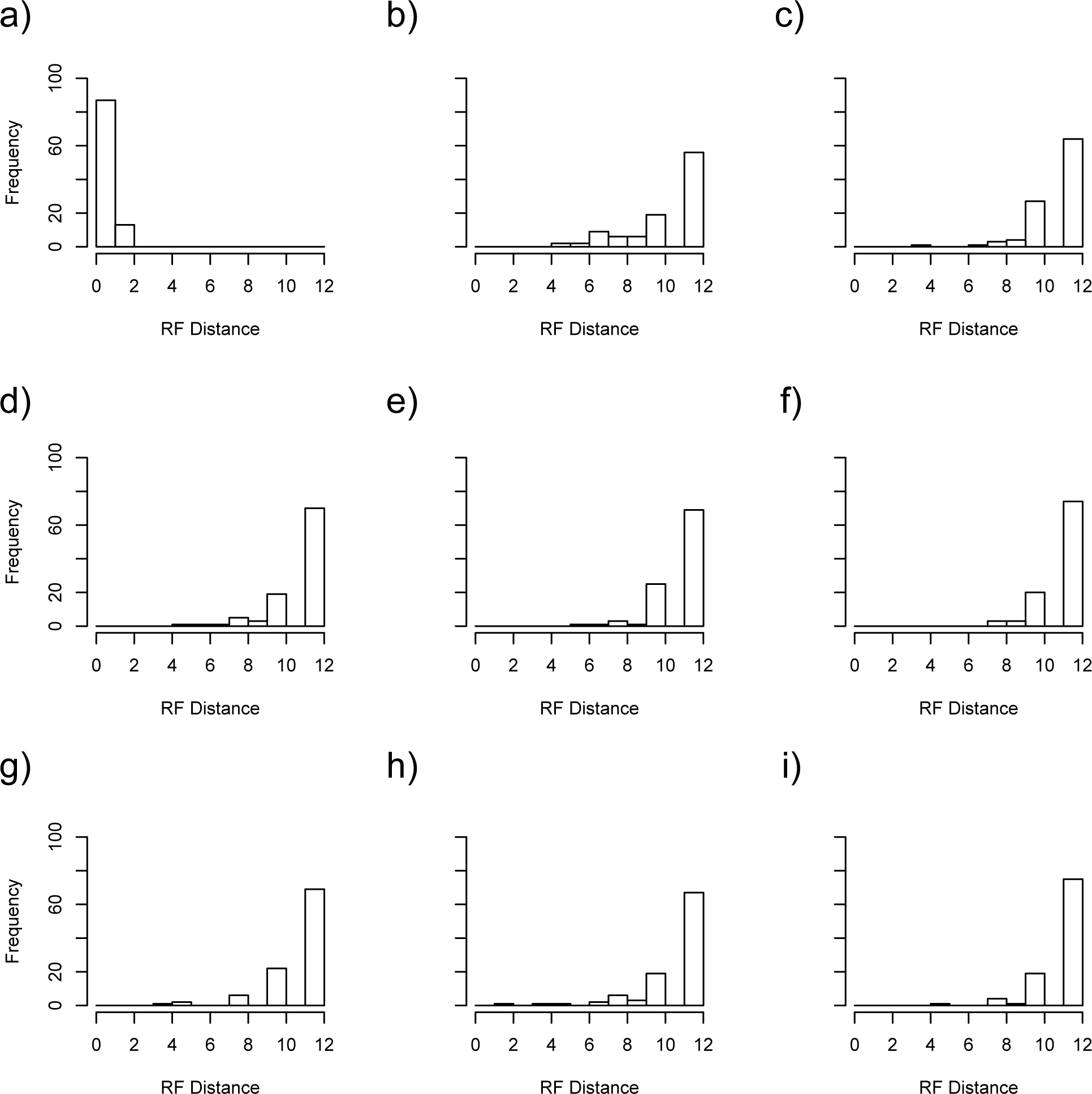
Frequency histogram showing the distribution of Robinson-Foulds distances between estimates using STEM and the correct tree under settings: a) D1, b) D2, c) D3, d) D4, e) D5, f) D6, g) D7, h) D8, and i) D9.

**Figure S6:**
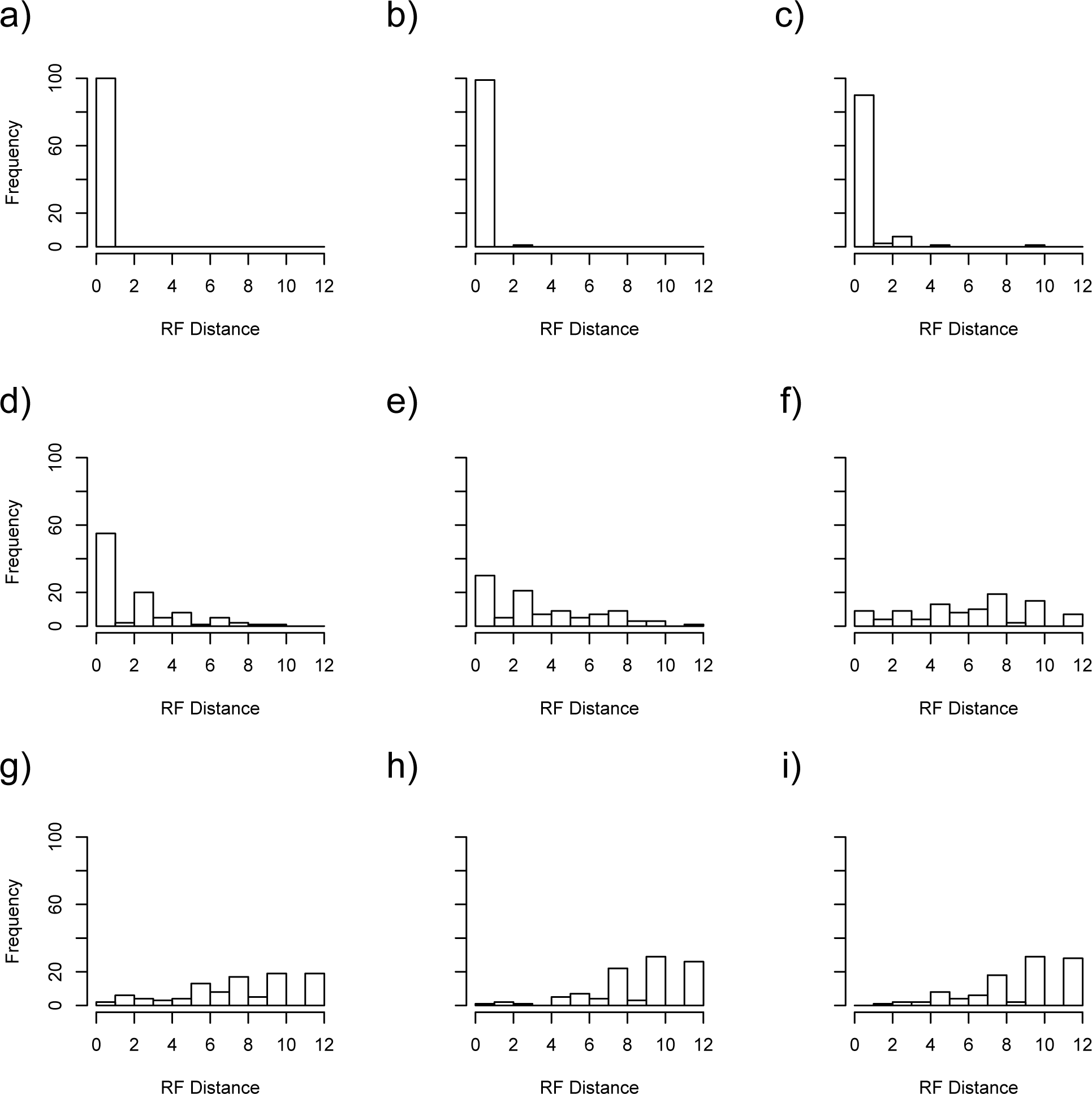
Frequency histogram showing the distribution of Robinson-Foulds distances between estimates using STEST (M0) and the correct tree under settings: a) D1, b) D2, c) D3, d) D4, e) D5, f) D6, g) D7, h) D8, and i) D9.

**Figure S7:**
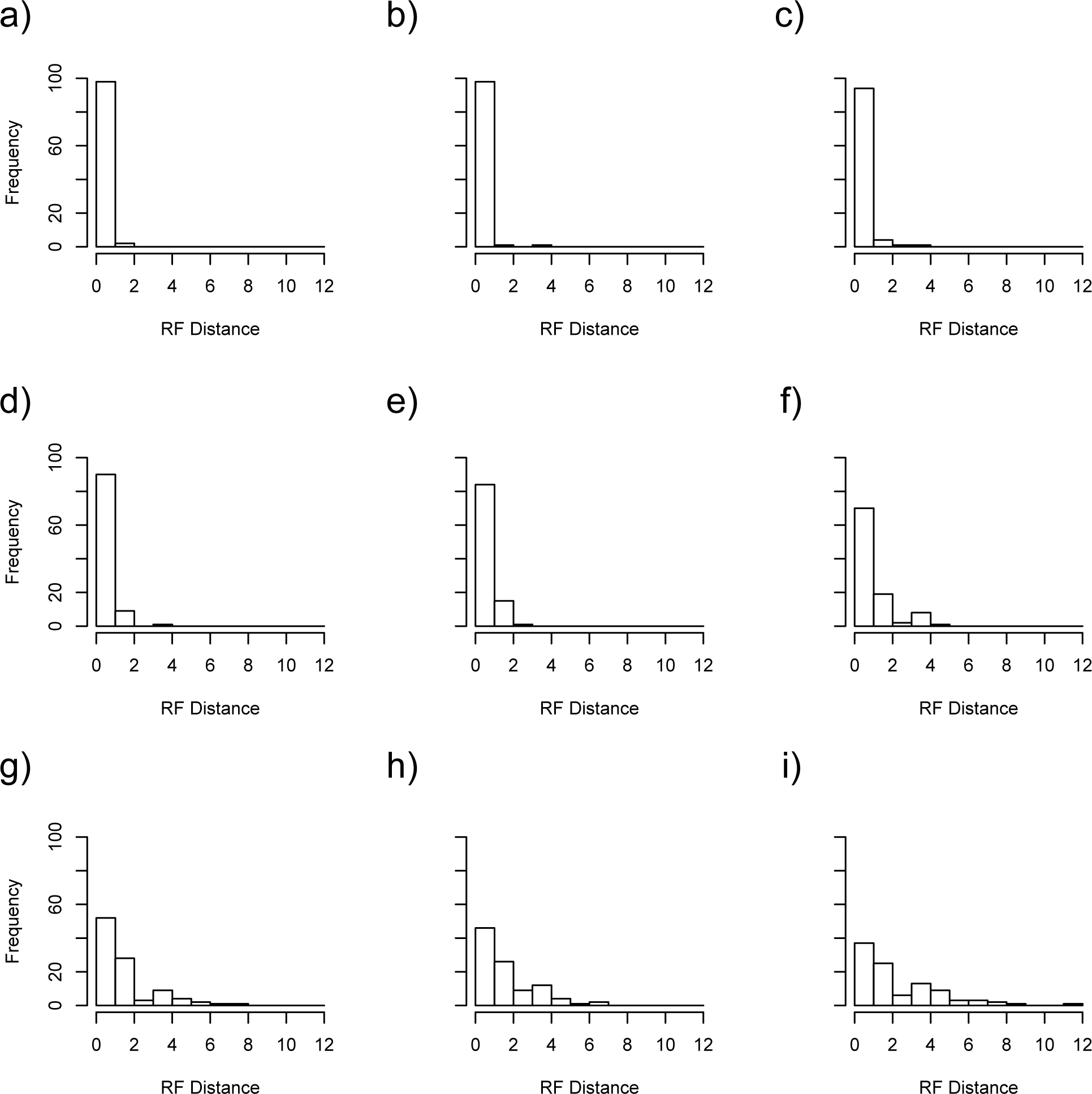
Frequency histogram showing the distribution of Robinson-Foulds distances between estimates using STEST (M1) and the correct tree under settings: a) D1, b) D2, c) D3, d) D4, e) D5, f) D6, g) D7, h) D8, and i) D9.

**Figure S8:**
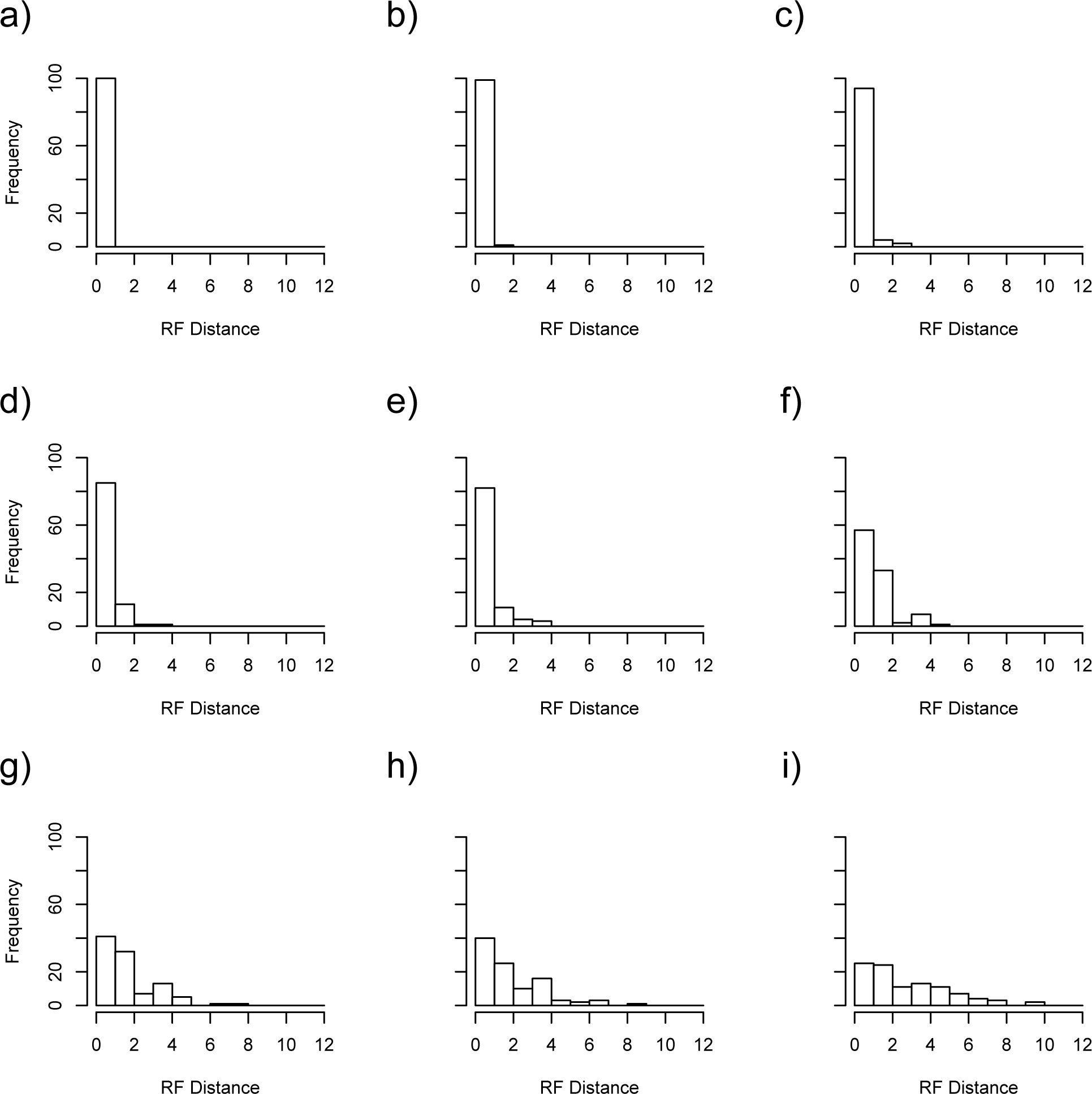
Frequency histogram showing the distribution of Robinson-Foulds distances between estimates using STEST (SIM3s) and the correct tree under settings: a) D1, b) D2, c) D3, d) D4, e) D5, f) D6, g) D7, h) D8, and i) D9.

### Empirical Study Results

The species tree estimates obtained using STEST (M0) and STEST (M1) both agree with the species tree topology obtained by Rannala and Yang (2003), which is (((H,C),G),O). The speciation time estimates 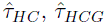 and 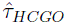 are listed in Table 7 in units of expected number of mutations per site, as in Rannala and Yang (2003). The running times are 14 seconds and 95 seconds for STEST (M0) and STEST (M1), respectively, on a Linux machine with two eight core Xeon E5-2680 (2.8 GHz) CPUs and 384 GB ram.

**Table 7:** Speciation time estimates for *τ*_1_, *τ*_2_ and *τ*_3_.

| | $\hat{\tau}_{HC}$ | $\hat{\tau}_{HCG}$ | $\hat{\tau}_{HCGO}$ |
| --- | --- | --- | --- |
| STEST (M0) | 0.00384 | 0.00571 | 0.01279 |
| STEST (M1) | 0.00396 | 0.00572 | 0.01331 |
| STEST (SIM3s) | 0.00384 | 0.00569 | $> 0.0223$ |

When attempting to use STEST (SIM3s) to estimate the species tree, we found that the SIM3s method was not able to estimate the speciation time when both taxa R (the outgroup) and O were included. Thus Table 7 gives the speciation time estimates for the other divergences, and only a lower bound on the speciation time for the split between taxa H, C, G and O. The total time to carry out this analysis was 130 seconds on the same machine.

## Discussion and Conclusion

### Simulation Study 1

When gene flow does not exist, STEST (M1) and STEST (SIM3s) perform worse than *BEAST and STEST (M0) in the short speciation interval scenario (Fig. 8a), which can be explained by the fact that short *τ*_1_ causes many deep coalescent events. The methods M1 and SIM3s may mistakenly attribute the species tree-gene tree conflicts partially to gene flow. This is possible because M1 and SIM3s only model the in-population processes within two focal populations (e.g., when *τ*_1_ is estimated, population 1 and population 2 are the two focal populations) and ignore the migration of any alleles between the two focal populations and any other populations (e.g., one allele is moved to population 123 at the time interval *τ*_2_, see Fig. 6). In the long speciation interval scenario, *τ*_1_ = 2 is long enough, which implies that there are not so many deep coalescent events. All the methods have similar good performance (estimation accuracy all above 94%, Fig. 8b). This also demonstrates the influence of deep coalescence in phylogenetic inference problems.

When gene flow does exist, *BEAST and STEST (M0) perform excellently in the short speciation interval scenarios (estimation accuracy above 86%, see Fig. 8a). This is because they assume the gene tree species tree conflicts are exclusively due to deep coalescence and short speciation interval causes many deep coalescent events, which exert much more influence on these conflicts than gene flow. Their performance decreases as the migration rate increases because the incongruence between gene trees and the species tree is influenced more and more by gene flow. The reason why STEST (M1) and STEST (SIM3s) do not perform as well as *BEAST and STEST (M0) might be the same as in the cases when there is no gene flow, i.e., gene flow occurs between two focal populations and the other populations. There are two possible cases in the presence of gene flow: the first is that before time *τ*_1_, gene flow occurs between the two focal populations and other populations in both directions, the second is that at time *τ*_2_, one linage is moved from an out population into the focal populations. The difference in STEST (M1) and STEST (SIM3s)’s performance may be due to either M1 and SIM3s’ different ability to estimate speciation times, or their different tolerance to the violation of their assumptions in our approach. Further study can be designed to find out which reason is more plausible. For now, we only want to evaluate the performance of the idea to estimate species trees. STEM’s performance curve is different. It decreases dramatically as migration rate increases. The reason is that as gene flow increases, the minimal coalescent time tends to zero. Therefore, STEM produces a lot of unresolved species trees, which implies that the data doesn’t have enough information for species tree estimation through STEM’s approach.

In the long speciation interval scenario, deep coalescence is no longer a problem. The incongruence between gene trees and species trees is mostly due to gene flow. Therefore, STEST (M1) and STEST (SIM3s) outperform *BEAST, STEST (M0) and STEM almost everywhere. Different from the short speciation interval scenarios, STEST (M1) and STEST (SIM3s)’s performance curves are very similar to each other, which indicates that their difference in performance is related to the many deep coalescent events in the previous scenarios. In most of the cases, *BEAST performs better than STEST (M0), which performs better than STEM. The performance curves are also decreasing as the migration rate increases. However, the slope of the performance curve is steeper than that in the short speciation interval scenario. The possible reason is that longer speciation interval allows more migration events when the migration rates are the same. Therefore, in the long speciation interval scenario, the same amount of increment in migration rate produces a larger increase in the number of migration events, which makes the performance of these methods decrease more.

There are multiple possible reasons why the performance of STEST (M1) and STEST (SIM3s) decrease when the migration rate increases. The first one is that the assumption that no other populations are exchanging genes with the focal populations in the M1 model and SIM3s model is violated. When there are not so many migration events, such violation does not matter a lot. However, when the speciation interval is long and migration rate between the focal populations and the unfocal populations is large, these methods are no longer applicable to estimate speciation times between two species. The second possible reason is that when the migration rate is large, the likelihood surface becomes bizarre. Therefore it is more difficult to locate the global maximum of the likelihood function. To improve this, M1 and SIM3s could be replaced by better methods (if any were developed) to estimate speciation times with the presence of gene flow.

### Simulation Study 2

When gene flow does not exist, all methods perform well in the cases when *τ*_1_ = 1 is moderate. When *τ*_1_ = 2 is long, STEST (M0) and STEST (M1) still perform very well in terms of estimation accuracy (*>* 95%). However, STEST (SIM3s) and STEM’s estimation accuracy fall slightly below 80%, which is different from the first simulation study, in which case all methods have good performance. One possible reason is that *τ*_2_ *- τ*_1_ = 1 in this case and *τ*_2_ *- τ*_1_ = 2 in the first simulation study. Thus, long ancestral speciation intervals might be helpful in species tree estimation in the presence of gene flow. Another possible reason is that larger trees are more difficult to estimate.

When gene flow does exist and *τ*_1_ = 1 is moderate, STEST (M1) performs the best when the migration rate is small (*<* 0.75), which implies deep coalescence is the main reason for the gene tree-species tree conflict. As expected, STEST (M1)’s performance starts to fall behind STEST (M1) and STEST (SIM3s) when the migration rate is large enough, which means that gene flow becomes the overwhelming factor for the conflict. STEST (M1) again performs consistently better than STEST (SIM3s). This could be the same reason as in simulation study 1. STEM again performs the best. This could also be explained by the same reason as in simulation study 1.

When *τ*_1_ = 2 is large, STEST (M1) outperforms all the other methods since more migration events are allowed even when migration rate is small. STEST (SIM3s) performs worse than STEST (M1) when the migration rate is smaller than 0.10. This might have the same reason as in the *τ*_1_ = 1 cases. When the migration rate is larger than 0.10, STEST (SIM3s) and STEST (M1) have similar performance. Their accuracy curves decrease more dramtically than the *τ*_1_ = 1 cases, because larger *τ*_1_ allows more migration events for the same increase in migration rate. It also decreases more dramtically than the long speciation interval scenarios in simulation study 1. This is because in simulation study 1, only one population exchanges genes with the two focal populations, while in this case, there are 5 more populations exchanging genes with the two focal populations before *τ*_1_. STEST (M0)’s performance decreases much more dramatically than in the previous cases, which shows it cannot deal with data subject to large migration rates. When migration rates are very large, speciation boundaries are not clear, and thus this behavior is not unexpected.

### Empirical Study

In the empirical study, all of the STEST-based methods STEST (M0), STEST (M1) and STEST (SIM3s) yield the correct tree topology within three minutes. The rapid and accurate performance of these methods for these data demonstrates the potential for the application of these methods to large-scale empirical data.

### Conclusion

To summarize, STEST (M0) provides an alternative approach to *BEAST for estimation of species trees in the presence of deep coalescence. It is much faster and has a comparable estimation accuracy. When the data follow the n-island migration model, STEST (M0) is appropriate for species tree estimation when the speciation interval for migration is short. When the speciation interval for migration is moderate, STEST (M0) is recommended for data subject to small migration rates and STEST (M1) is recommended for data subject to large migration rates. When the speciation interval for migration is long, STEST (M1) is the better choice.

There are multiple ways to improve the performance of our methods. One way, for example, is to develop better speciation time estimation methods. Our idea is to use the speciation time estimates as distances to estimate species trees. The better quality the speciation time estimates are, the better accuracy our method will have. Another way is to find the best strategy to accommodate different and more informative data types. For example, extension of the SIM3s, M0 and M1 methods to handle multiple sampled lineages per species would allow our method to be applied in this setting.

